# Maize AFP1 confers antifungal activity by inhibiting chitin deacetylases from a broad range of fungi

**DOI:** 10.1101/2021.11.02.466889

**Authors:** Lay-Sun Ma, Wei-Lun Tsai, Raviraj M. Kalunke, Meng-Yun Xu, Yu-Han Lin, Florensia Ariani Damei, Hui-Chun Lee

## Abstract

Adapted plant pathogenic fungi deacetylate chitin to chitosan to avoid host perception and disarm the chitin-triggered plant immunity. Whether plants have evolved factors to counteract this fungal evasion mechanism in the plant-pathogen interface remains obscure. Here, we decipher the underlying mechanism of maize cysteine-rich receptor-like secreted proteins (CRRSPs)-AFP1, which exhibits mannose-binding dependent antifungal activity. AFP1 initials the action by binding to specific sites on the surface of yeast-like cells, filaments, and germinated spores of the biotrophic fungi *Ustilago maydis*. This could result in fungal cell growth and cell budding inhibition, delaying spore germination and subsequently reducing fungal viability in a mannose-binding dependence manner. The antifungal activity of AFP1 is conferred by its interaction with the PMT-dependent mannosylated chitin deacetylases (CDAs) and interfering with the conversion of chitin. Our finding that AFP1 targets CDAs from pathogenic fungi and nonpathogenic budding yeast suggests a potential application of the CRRSP in combating fungal diseases and reducing threats posed by the fungal kingdom.

## Introduction

Plants safeguard the apoplast environment by deploying cell-surface localized pattern recognition receptors (PRRs) to sense potential danger signals via recognition of pathogen-associated molecular patterns (PAMP) or plant-derived damaged-associated molecular patterns (DAMP)^1, 2^. This alerts the plant immune system and leads to the delivery of a mix of plant defense molecules to the apoplast to ward off pathogenic intruders^3^. Adapted pathogens are able to breach this defense barrier by shielding their cell surfaces, sequestering PAMPs, and modifying cell-surface glycans to avoid recognition and escape from the host attack at the initial steps of pathogenesis^4^. In return, plants evolve new weapons to counteract the fungal camouflage strategies.

The biotrophic fungus *Ustilago maydis* causes smut disease in maize by inducing large tumors in which the fungal hyphae proliferate and form spores^5^. The complex interplay between *U. maydis* and maize at the interface is mainly governed by an arsenal of fungal effector proteins induced in consecutive waves after the fungus contacts and colonizes maize^6^. To date, a few functionally characterized effectors have been reported to act in the apoplasts to suppress plant immunity and to protect hyphae from the attack of plant defense molecules. *U. maydis* Fly1 prevents the release of chitin by destabilizing plant chitinases^7^, Rsp3 shields fungal hyphae to block the antifungal activity of maize AFP1 proteins^8^, and Pit2 and Pep1 inhibit the activities of plant cysteine protease and peroxidases respectively^9,10^. *U. maydis* also possesses six active chitin deacetylases to convert chitin to chitosan to sustain fungal cell viability and evade recognition by chitin receptors and promote virulence^11^.

Plant cysteine-rich receptor-like kinase (CRK) is one type of the transmembrane PRR proteins featured with domains of unknown function 26 (DUF26; stress-antifungal domain PF01657) which harbors a conserved cysteine motif (C-X8-C-X2-C) in their ectodomain^12, 13^. CRKs have been implicated in plant defense, oxidative/salt stress and salicylic acid responses^14–17^. DUF26 domain has also been found in some plasmodesmata-localized proteins (PDLPs) and cysteine-rich receptor-like secreted proteins (CRRSPs)^12, 13^. PDLPs have a configuration similar to CRKs but lack a kinase domain. They regulate plasmodesmata permeability and immunity and modulate callose deposition in plasmodesmata^17, 18^. CRRSPs harbor one or two copies of the DUF26 domain and are secreted to the apoplast upon pathogen infection^8, 19^. Two of the maize CRRSPs (AFP1 and AFP2), display mannose-binding-dependent antifungal activity and are significantly upregulated upon the perception of *U. maydis*^8^. AFP1/2-silenced plants are more susceptible to the infection by *U. maydis* and *Colletotrichum graminicola*^8^, suggesting that maize AFP1/2 are involved in plant defense against a broad range of fungal pathogens. So far, the antifungal activities of CRRSPs have been demonstrated for maize AFP1^8^ and *Ginkgo biloba* Gnk2^20^ but not for cotton CRR1 which was shown to stabilize plant chitinases ^21^. Despite the importance of the DUF26-containing proteins in regulating biotic and abiotic stress responses, their biological function in plant immunity is not clear, and the antifungal mechanism of the CRRSP has not yet been elucidated.

In this work, we study maize AFP1 to provide mechanistic insights into the mode of action of plant CRRSP. The provided evidence supports that maize AFP1 proteins could inhibit fungal growth by interfering with the activity of chitin deacetylases which are essential for fungal development and virulence. Interfering with the alteration of a fungal cell wall important component highlights the potential use of CRRSPs as antifungal agents to act against a broad-range of pathogenic and non-pathogenic fungi.

## Results

### AFP1 localizes on the growing tips and bud necks of yeast-like cells in *Ustilago maydis*

AFP1 displays a mannose-binding dependent antifungal activity^8^. However, the underlying molecular mechanism has not yet been deciphered. To investigate the initial action mode of AFP1, we examined the binding of AFP1 to *U. maydis* cells using immunolocalization. SG200 budding cells incubated with purified wild-type AFP1-His and mannose-binding-deficient AFP1*-His proteins (contains S34A, R115A, E126A, N144A, Q227A, and E238A mutations in the two mannose-binding sites)^8^ were subjected to immunostaining using the anti-His antibody and AF488-conjugated secondary antibody without any permeabilization and fixation (Fig. 1A). The fluorescence of AFP1-His was detected throughout the cell body and showed some accumulation in speckles on the cell periphery. Predominant fluorescence accumulation was seen at cell division zones and in the growing tips of emerging buds (Fig.1A, central panel and enlarged panel on the right). Besides that, AFP1 fluorescence was also detected at growing tips and on the surface of SG200 filaments induced by hydroxy-fatty acids on a hydrophobic surface without any permeabilization (Fig. S1A). AFP1*-His fluorescence could neither be detected on yeast-like cells nor on filaments of SG200 (Fig. 1A and Fig. S1A), suggesting that the AFP1 binding is dependent on its mannose-binding ability.

**Fig. 1.**
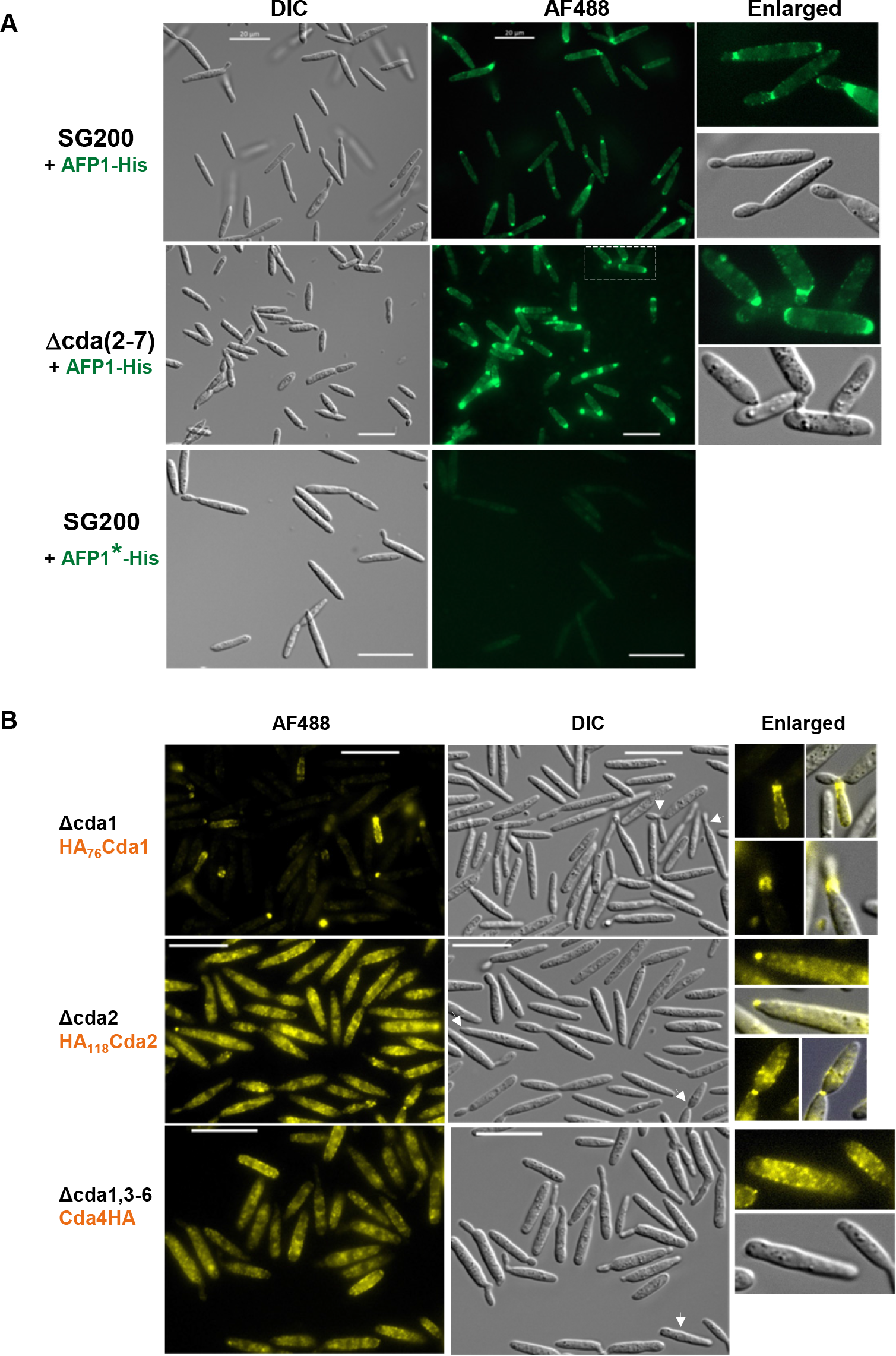
AFP1 and *U. maydis* CDAs display similar localization patterns. (A) Yeast-like cells of indicated strains were incubated with either AFP1-His or AFP1*-His (defects in mannose-binding) proteins and followed by immunostaining with an anti-His antibody and an AF488-conjugated secondary antibody. Bars, 20 μm. (B) Cells expressing HA-tagged CDA proteins under constitutive promoter *otef* in indicated deletion strains were subjected to immunostaining using anti-HA antibody and an AF488-conjugated secondary antibody to localize CDA proteins. White arrows indicate cells enlarged in right panels. Bars, 20 μm.

### *U. maydis* CDAs display similar localization patterns as the maize AFP1 proteins

The localization pattern of AFP1 in *U. maydis* cells is similar to that of *U. maydis* chitin synthases (CHS) and chitinases (CTS), and is also correlated with the sites in the cell wall where chitin and chitosan are accessible to antibodies in yeast-like cells^11, 22–24^. This prompted us to examine if AFP1 target the fungal CHS or CTS proteins. To test this idea, we examined the interaction of AFP1 with either CHS or CTS using a yeast-two hybrid (Y2H) assay. However, we could not detect the interactions between AFP1 and CHS or CTS in the Y2H analysis (Fig. S2).

The conversion of chitin to chitosan is important for fungal development and is one of the fungal evading strategies to avoid recognition by host plants. Chitin deacetylases might co-localize with CHS to access to nascent chitin substrates. We posit that AFP1 might target chitin deacetylases (CDAs) to block the deacetylation of chitin. The *U. maydis* genome contains seven *cda* genes that encode six C-terminal GPI (glycosylphosphatidylinositol) anchored CDAs and one secreted CDA protein (Cda4). While *cda6* is a pseudogene and not expressed^11^, all active *cda* genes were differentially expressed in the axenic culture of SG200 cells (Fig. S3). Since the deletion of all *cda* genes is lethal to *U. maydis*, we initially examined the AFP1-His localization in a *cda* sextuple deletion mutant (Δcda2-7) that lacks six out of the seven *cda* genes^11^. Surprisingly, instead of detecting a weak AFP1 accumulation, we observed a more intense AFP1-His fluorescence in the Δcda2-7 mutant comparing to the parental SG200 cells (Fig. 1A). Since a significant increase in the expression of the *cda1* gene was detected in the deletion mutants Δcda2-6 and Δcda2-5 (Yanina S. Rizzi, personal communication), the *cda1* gene expression might also be upregulated in Δcda2-7 to compensate for the loss of other *cda* genes. This prompts us to speculate that the remaining Cda1 could be a potential binding target of AFP1.

Since the localization of CDA proteins in *U. maydis* has not been reported, we next explored the immunolocalization of the two highly expressed Cda1 and Cda2 proteins in *U. maydis* cells. We generated an N-terminal HA-tagged Cda1 protein (HACda1). However, we failed to immunolocalize HACda1 proteins on the cell surface without permeabilization even though the protein was detected in cell extracts (Fig. S4), suggesting that the N-terminus of Cda1 protein might undergo processing before it becomes membrane-anchored. An HA-tag was then inserted at the amino acid 76 of Cda1 (HA76Cda1), and overexpressed in the *cda* quintuple mutant (Δcda1,3-6)^11^ and *cda1* deletion mutant (Δcda1) to avoid the interference of endogenous Cda1 to compete with HA76Cda1 for the localization. In both backgrounds, HA76Cda1 fluorescence appeared in speckles at the growing tips and around the cell periphery of emerging cells but rarely accumulated in the cell periphery of mother cells (Fig. 1B and Fig. S1B). Occasionally, we also visualized the HA76Cda1 speckles on the cell division zones (Fig. 1B; Enlarged panels). In filaments of Δcda(1,3-6)-HA76Cda1 strain, the HA76Cda1 fluorescence was also detected on the surface and growing tips of filamentous cells (Fig. S1B), which was similar to the localization of AFP1-His on filaments of SG200 and Δcda(1,3-6)-HA76Cda1 (Fig. S1A and S1C). In comparison to the HA76Cda1 localization pattern, HA118Cda2 speckles mainly appeared in the cell periphery of the *cda2* deletion mutant that overexpressed HA118Cda2 proteins (Δcda2-HA118Cda2) (Figure 1B). Additionally, few HA118Cda2 speckles appeared at the cell tips and cell division zones (Fig. 1B; Enlarged panels). Further localization analysis of the secreted Cda4 proteins expressed in Δcda(1,3-6)-Cda4HA strain also revealed that the Cda4HA proteins exclusively localized to the cell periphery (Fig. 1B). These results show that the CDAs display similar localization patterns as the AFP1 in yeast-like cells and filaments of *U. maydis* cells.

### Maize AFP1 colocalizes and interacts with *U. maydis* Cda1

We next examined the possible interaction between AFP1 and Cda1 by analyzing their co-localization. Δcda(1,3-6)-HA76Cda1 cells were subject to immunostaining before being incubated with the fluorescent probe DyLight550-labeled AFP1-His proteins (AFP1-His^550^). We found that the fluorescence of HA76-Cda1 and AFP1-His^550^ overlapped at bud necks and in the growing tips of the cells (Fig. 2A). To examine the interaction between AFP1 and Cda1 in Δcda (1,3-6)-HA76Cda1 cells at suboptical resolutions, we measured acceptor photobleaching FRET (Fluorescence resonance energy transfer) efficiency^25–27^ by bleaching the acceptor - AFP1-His^550^ to eliminate or reduce energy transfer from the donor HA76Cda1, and thereby yielding an increase in the donor fluorescence (Fig. 2B). The FRET efficiency of HA76Cda1 after bleaching AFP1-His^550^ increased significantly with an average of 23%, compared to 4% in the negative control (WGA-AF488, wheat germ agglutinin conjugated to Alexa-Fluor 488 that recognizes chitin). This result suggests that Cda1 could be an AFP1-interacting target in the *U. maydis*.

**Fig. 2.**
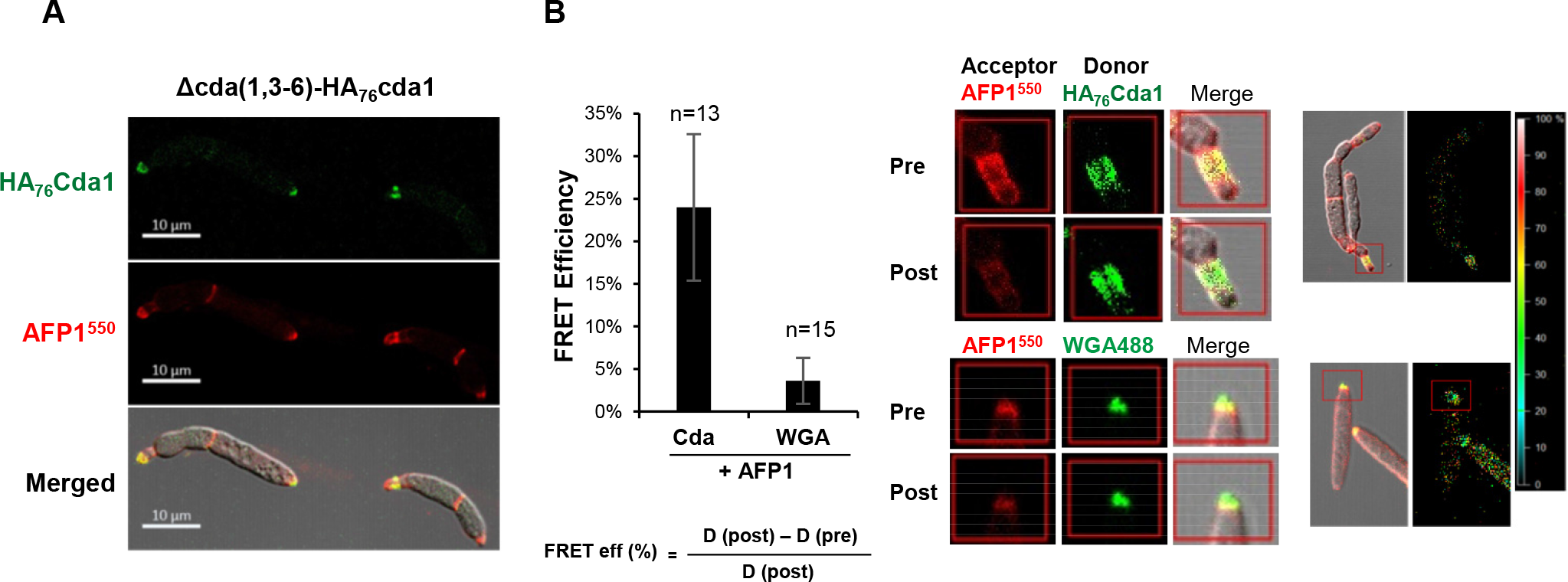
The co-localization of Cda1 and AFP1. Cells of Δcda(1,3-6)-HA_76_cda1 strain were immunostained to localize HA_76_cda1 (A, B) or stained with WGA-AF488 to locate chitin (B) before incubation with DyLight 550 labeled AFP1-His proteins (AFP1^550^) and visualized the fluorescence by confocal microscopy. (B) FRET efficiency was calculated using the indicated equation. Number of spots from three biological replicates used in the analysis as indicated above column. Values represent mean ± standard deviation. Representative images of cells showing fluorescence of AFP1^550^ and HA_76_Cda1/WGA-AF488 were taken and measured before and after the photobleaching of acceptor AFP1^550^ (middle panel). Cell images with overlapped fluorescence of HA_76_Cda1 or WGA-AF488 and AFP1^550^ and the signal intensity in overlapping regions are shown (right panel).

### AFP1 targets more than one *U. maydis* chitin deacetylases and reduces CDA activity

The interaction of AFP1 with Cda1 prompted us to investigate whether AFP1 also directly interacts with other *U. maydis* CDA proteins. Using yeast-two hybrid (Y2H) assay, we found that AFP1 interacted with Cda1, 3, 5, and 7 but not with Cda2 and Cda4 even though the proteins expressed at similar levels (Fig. 3A and Fig. S5A). When adding a competitive inhibitor 3-AT (3-amino-1, 2, 4-triazole) to increase the stringency of the selection, Cda1 and Cda7 were the only two targets left in the interaction and AFP1 displayed a stronger interaction with the Cda1 (Fig. S5B). Furthermore, the CDA domain of Cda1 was sufficient to mediate the interaction with AFP1 in the Y2H (Fig. S5C).

**Fig. 3.**
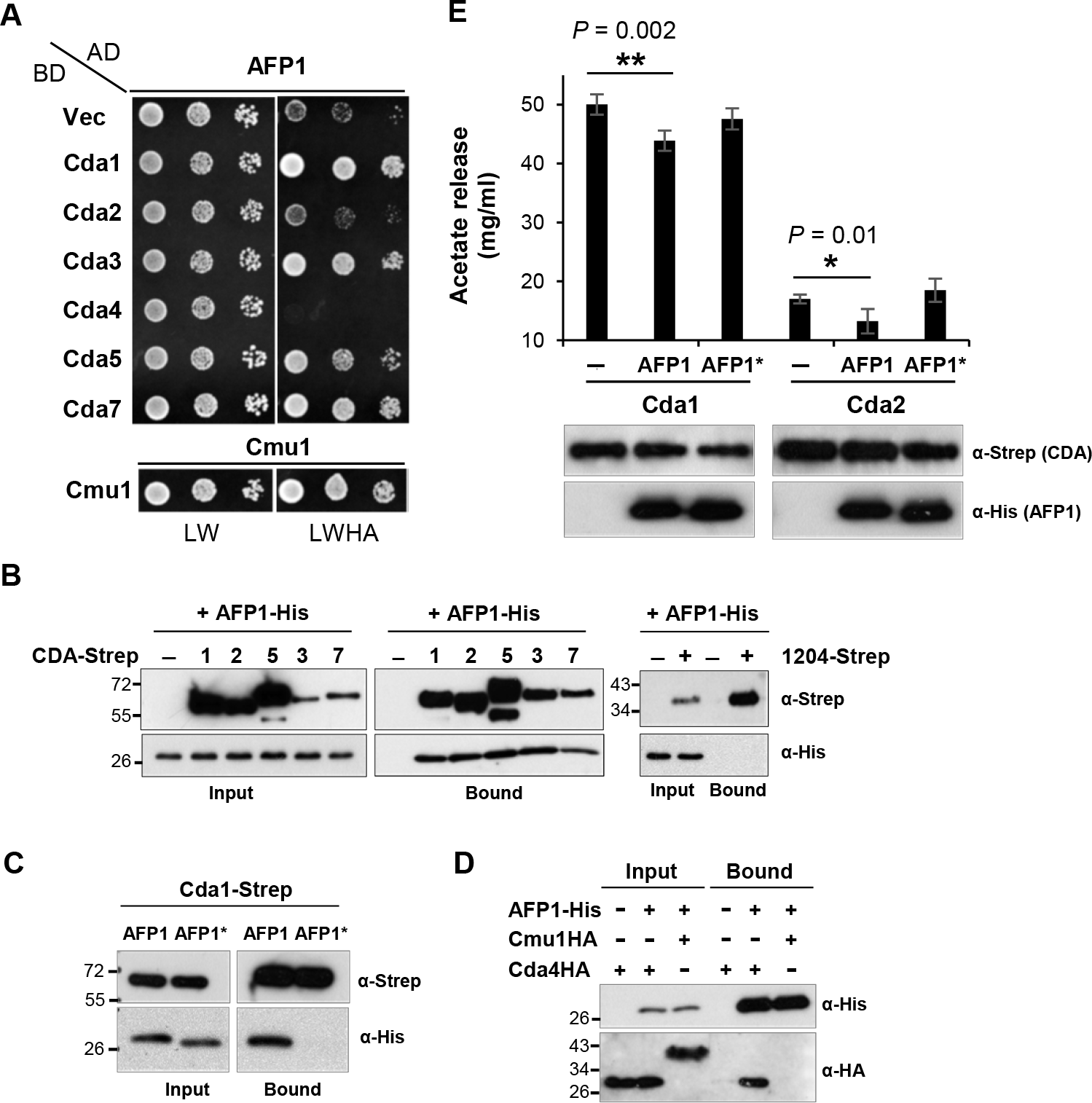
AFP1 acts on CDA proteins to interfere chitin deacetylase activity. (A) Yeast two-hybrid assay to detect interaction of AFP1 and CDA proteins. Yeast cells with indicated plasmids were grown on SD/-Leu/-Trp (LW) and SD/-Leu/-Trp/-His/-Ade (LWHA) plates for 3 days. Self-interaction of chorismate mutase (Cmu1) was served as the positive control and AD-AFP1/BD as negative control. Similar results were observed in at least two independent experiments. (B-C) In vitro pull down assay of CDA-Strep and AFP1 proteins. Indicated C-terminal Strep-tagged CDA or 1204 (*U. maydis* secreted proteins; unrelated protein control) proteins were immobilized on Strep-Tactin agarose beads and served as baits to pull down AFP1-His or AFP1*-His proteins. (D) AFP1-His proteins were immobilized on Ni-NTA agarose beads and incubated with C-terminal HA-tagged Cda4 and Cmu1 (negative control). (E) Enzyme activity of Cda1 and Cda2 in presence of AFP1-His and AFP1*-His. Cda1-Strep and Cda2-Strep proteins immobilized on Strep-Tactin agarose beads were treated with or without AFP1 proteins before incubated with GlcNAc_5_ (A5) substrates. Data given represent the mean ± SD values of the results from three independent experiments. Immunoblots showing similar amount of proteins in each reaction and could be detected after incubation (lower panel). Asterisks indicate significant differences of CDA activity in respective reaction compared with buffer control (-) determined by a two-tailed Student’s *t*-test. *P* values are shown.

The AFP1-CDA interaction was further validated by immobilizing secreted C-terminal Strep-tagged Cda1, 2, 3, 5, and 7 deleting respective GPI-anchor proteins (CDA-Strep) on Strep-Tactin agarose beads and followed by incubation with purified AFP1 proteins. Unexpectedly, the wild-type AFP1-His proteins were pulled down by all CDA-Strep proteins but not by an unrelated protein control-1204-Strep (UMAG01204, an *U. maydis* signal-peptide containing protein) and the buffer control (Fig. 3B). Notably, the Cda1-AFP1 interaction also required the mannose-binding ability of AFP1 as the Cda1-Strep could not pull down AFP1*-His proteins (Fig. 3C). To analyze if AFP1 also interacts with the secreted Cda4 proteins lacking GPI anchor, secreted Cda4HA (C-terminal HA-tagged Cda4) or Cmu1HA (*U. maydis* secreted chorismate mutase)^28^ from culture supernatants of strains Δcda(1,3-6)-Cda4HA or SG200-Cmu1HA was collected and incubated with the immobilized AFP1-His proteins on Ni-NTA agarose beads. AFP1-His proteins could pull down Cda4HA but not Cmu1HA proteins, and non-specific binding of Cda4HA proteins to the beads was also not detected (Fig. 3D). These results indicate that AFP1 could interact with all six *U. maydis* CDA proteins. Since the CDA proteins share a conserved NodB homolog domain^11, 29^, AFP1 likely recognizes the conserved structural fold of CDA proteins.

We next studied the impact of AFP1-CDA interaction on the chitin deacetylase activity by measuring the release of acetate. The Cda1 or Cda2-Strep proteins were immobilized on beads and incubated with either AFP1 proteins or buffer before adding the substrate GlcNAc5 (A5). The released acetate was significantly reduced when the Cda1-Strep and Cda2-Strep proteins were incubated with AFP1-His, but not when incubated with AFP1*-His or buffer control (Fig. 3E). Altogether, the results suggest that AFP1 binds to the CDA proteins and acts to reduce the deacetylase activity of CDA in a mannose-binding-dependent manner.

### Cda1 and Cda2 but not Cda4 are the substrates of PMT4

Considering AFP1-binding is the mannose-binding dependent, we reasoned that CDAs might be mannosylated. We thus analyzed the AFP1-binding in a mannosyltransferase deletion mutant of *U. maydis*, *pmt4*, which is defective in early infection-related development^30^. In the Δpmt4 strain, the prominent AFP1 fluorescence on the bud necks and scars almost diminished, whereas the intensity of speckles on the cell periphery became more evident (Fig. 4A and 4B). These results indicate that the CDAs localized at the cell division zone and the growing tips require the mannosylation activity of PMT4 and imply that the cell division localized Cda1 and Cda2 but not the cell-periphery localized Cda4 could be the targets of PMT4.

**Fig. 4.**
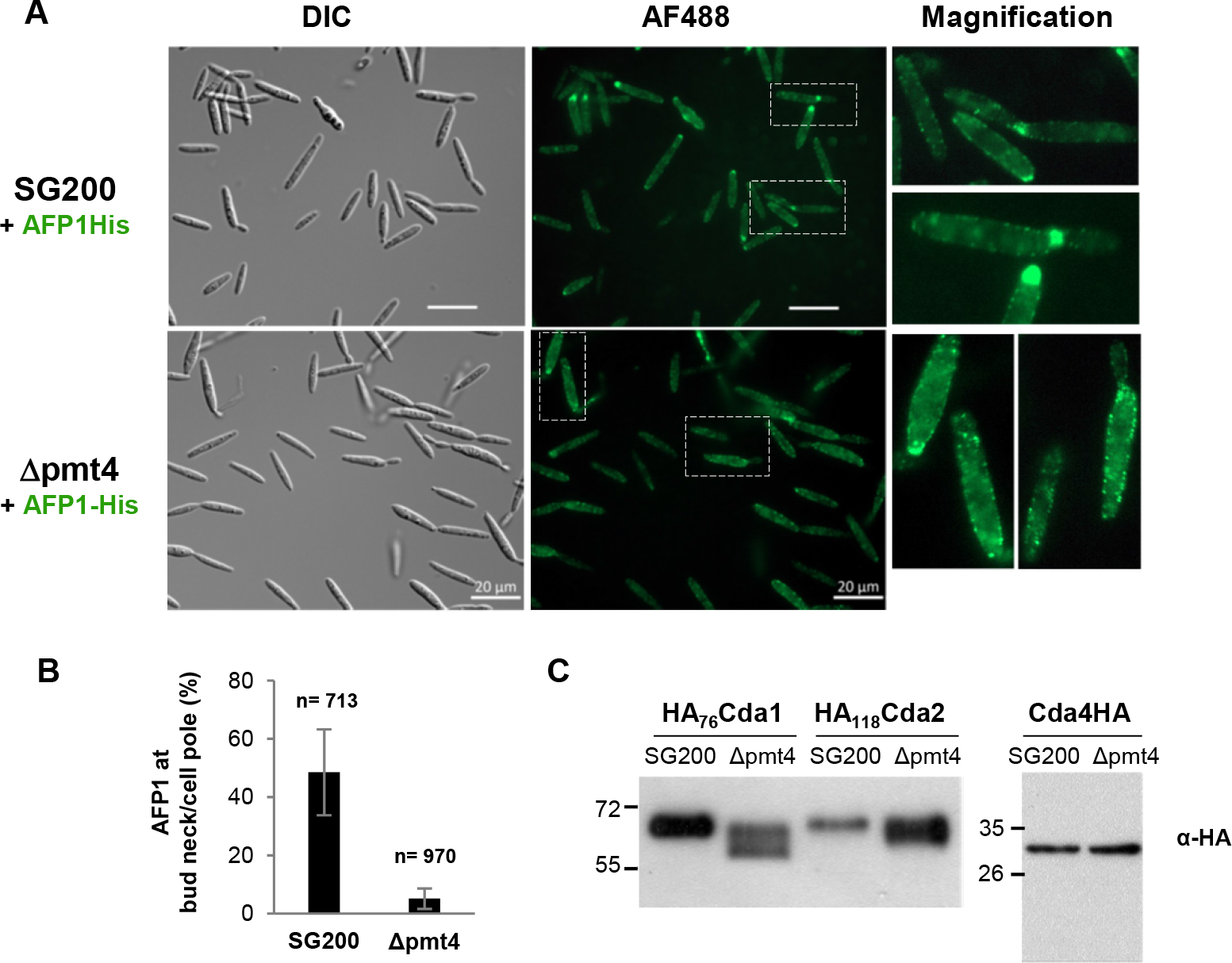
Pmt4 mannosylates Cda1 and Cda2. (A) Yeast-like cells of indicated strains were incubated with AFP1-His and followed by immunostaining with an anti-His antibody and an AF488-conjugated secondary antibody. Bars, 20 μm. Arrows indicate the bud neck/cell pole localization of AFP1. (B) Total number of cells (n) from three biological replicates in experiment a as indicated at above each column was analyzed. The percentage of cells with AFP1 binding at bud necks/cell poles was calculated. Values represent mean ± sd of three biological replicates. Arrows indicate AFP1 fluorescence on cell pole /bud neck of cells of Δpmt4 strain. (C) PMT4 mannosylates Cda1 and Cda2. HA_76_Cda1 and HA_118_Cda2 were overexpressed in either SG200 or Δpmt4 strains. The total protein extracts collected from the cell pellet of indicated strain were treated with (+) or without (-) deglycosylation enzyme mix (NEB Cat# P6039) according to the manufacturer’s protocol, and the glycosylation pattern of CDA proteins were analyzed by immunoblotting using anti-HA antibody. (D) Immunolocalization of HA_76_Cda and HA_118_Cda2 overexpressed in indicated strains. Bars, 20 μm.

We overexpressed HA76Cda1, HA118Cda2, and Cda4HA in SG200 and the *pmt4* deletion mutant strain and examined the migration patterns of the proteins in both backgrounds (Fig. 4C). Cda1 and Cda2 proteins displayed a smear and faster migration pattern when expressed in the *pmt4* mutant compared to the wild-type SG200 background (Fig. 4C). In contrast, the migration patterns of the cell-periphery localized Cda4HA in SG200 and the *pmt4* mutant background were similar. The result illustrates that Cda1 and Cda2 but not Cda4 are the substrates of PMT4. Further immunolocalization analysis of HA76Cda1 and HA118Cda2 revealed no difference in the localization patterns of the CDA proteins expressed in wild-type SG200 and the *pmt4* mutant (Fig. S6). Therefore, the deficiency in the PMT4-dependent mannosylation does not alter the localization of Cda1 and Cda2 but reduces the AFP1-binding. The incomplete abrogate AFP1-binding to cells of the *pmt* mutant suggests that either AFP1 binds to the additional CDA targets that are not mannosylated by PMT4, e.g., Cda4. Or, the decrease in the Cda1/Cda2 mannosylation may significantly reduce but not abolish the interaction of Cda1/2 and AFP1. The result links the mannose-binding of AFP1 to mannosylated CDAs.

### AFP1 binds and blocks spore germination by inhibiting cell budding and growth of fungi

Since spores are the likely propagules in which the infections occur in the field and the *cda* genes in *U. maydis* are upregulated and expressed during spore germination^11^, we hypothesized that AFP1 can bind *U. maydis* spores and inhibit its germination. To investigate this, we examined the AFP1 localization on germinated spores of *U. maydis*. Asynchronous germinated spores on agar were harvested and incubated with the AFP1 proteins before immunostaining. While background noises were found in cell debris and mucilage but not in promycelium of the AFP1*-His-treated spores (Fig. 5A), the AFP1-His fluorescence was seen at the bulged regions of spores which presumably are the sites of promycelium emergence (Fig. 5B) and the base of promycelium where it had protruded from spores (Fig. 5C-D). In addition, the fluorescence also appeared at the tips of promycelium (Fig. 5E-F) and the regions between two basidial cells (Fig. 5G-I). The results illustrate that AFP1 binds to germinating spores at distinct morphological stages during germination, as early as the promycelium first protrusion.

**Fig. 5.**
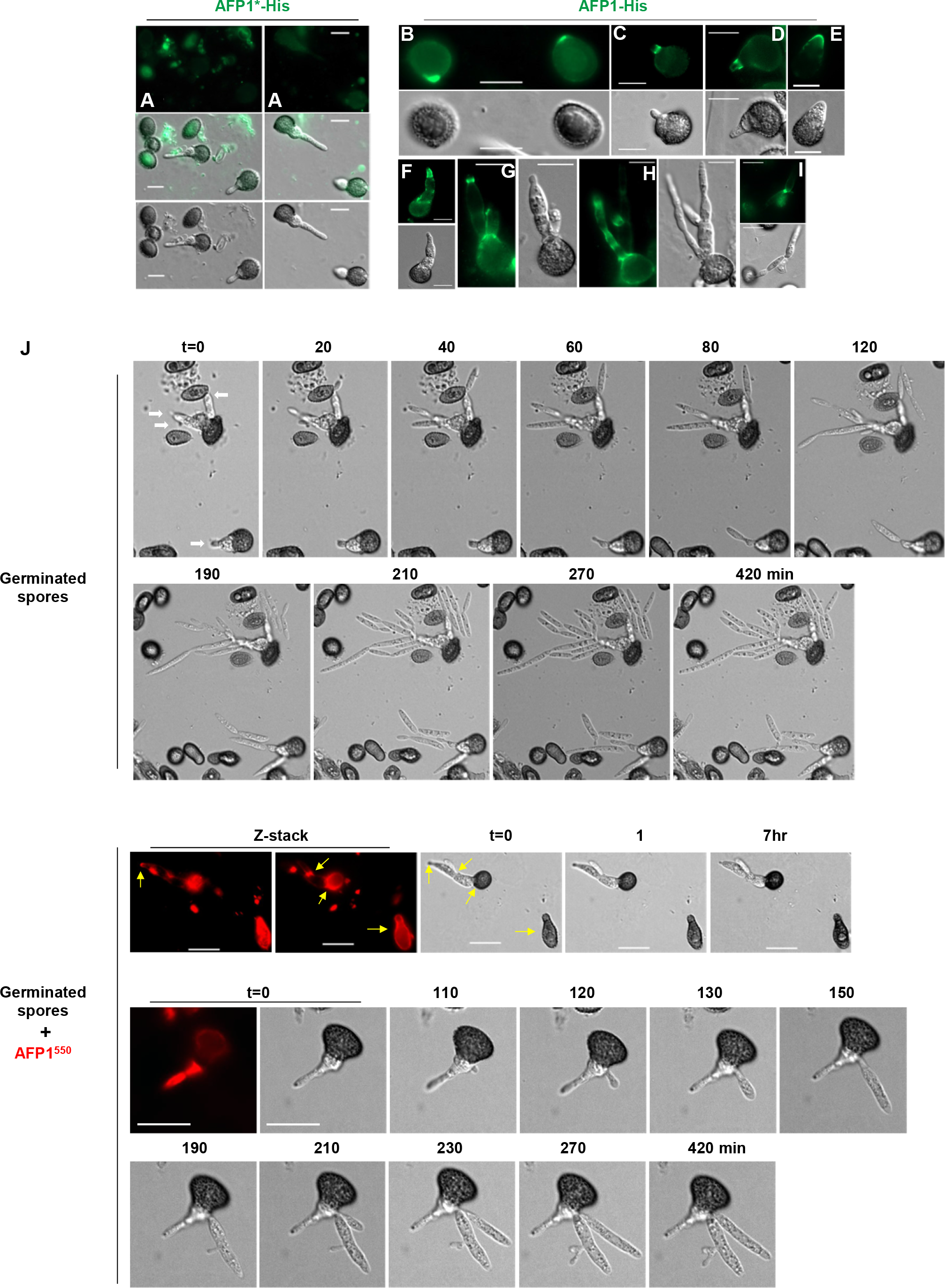
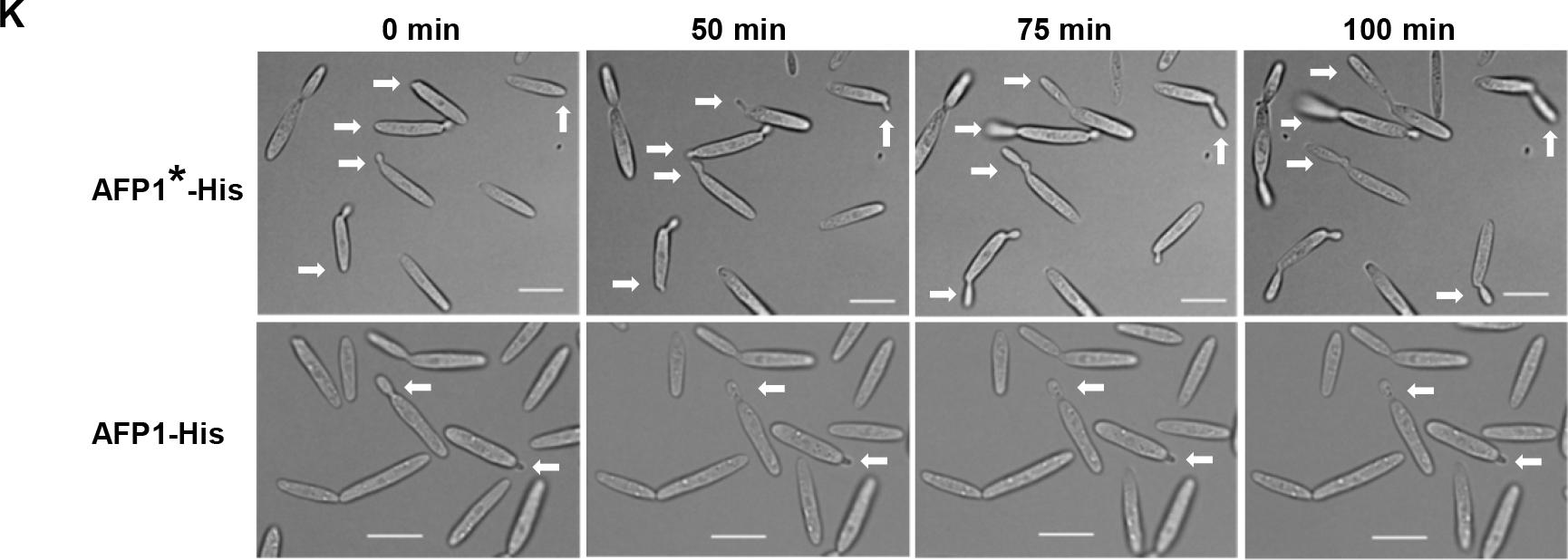
AFP1 inhibits spore germination, cell budding and the growth of budding cells. *U. maydis* FB1x FB2 germinated spores and SG200 yeast-like cells were treated with AFP1-His and AFP1*-His proteins in agar or liquid medium containing 2% nutrients. Bars,10 μm. (A-I) AFP1-treated germinated spores were immunostained with an anti-His antibody and an AF488-conjugated secondary antibody to visualize AFP1 fluorescence. j, Images of germinated spores with or without AFP1^550^ added were acquired at indicated time points. Yellow arrows in two Z-stack images indicate AFP1^550^-His binding sites on the germinated spore. (K) Images of AFP1-treated SG200 yeast-like cells were acquired at indicated time points. White arrows track the growth of cells over time in medium with AFP1-His or AFP1*-His proteins added.

Owing to asynchronous spore germination and low germination rate in *U. maydis*^31–34^, quantitative inhibitory assays of AFP1 on spore germination is challenging. To dissect the inhibitory activity of AFP1 on spore germination, we performed time*-*lapse live-cell imaging to monitor the germination process of individual spores in the presence or absence of DyLight550 labeled AFP1^550^-His proteins (Fig. 5J). In the absence of AFP1 proteins, most of the spores remained dormant and only a few promycelium emerged from spores, elongated, produced basidiospores, and then fell off. The germination process was slow down 5 hours later and was at a halt in 7 hours after nutrients depleted. The emergence and elongation of promycelium were detected between 20 minutes and 60 minutes, depending on the germination stages of spores (Fig. 5J and Fig. S7). Compared to germinated spores with no AFP1^550^ added, AFP1^550^-bound promycelium that showed stronger fluorescence signal was either completely stopped elongating (Fig. 5J; Upper panel) or the promycelium abandoned the AFP1^550^-bound basidial cells and produced a new basidial cell from another side to generate basidiospores and escaped the inhibitory effect of AFP1. However, this escaping process could delay germination by one to two hours (Fig. 5J; Lower panel). The results indicate that AFP1 binding could delay and block spore germination.

The inhibitory effect of AFP1 in yeast-like cells was also observed (Fig. 5K). Budding and growth of yeast-like cells were inhibited by the AFP1-His treatment but not by the AFP1*-His treatment where cells continued growing and budding. Therefore, this result reinforces the antifungal activity of AFP1 towards *U. maydis* yeast-like cells.

### AFP1 targets pathogenic and non-pathogenic fungi by acting on conserved CDAs

Having shown that AFP1 acts on the *U. maydis* CDAs to inhibit spore germination and cell growth, we next asked whether AFP1 also exerts inhibitory effects on other pathogenic and nonpathogenic fungi. We localized AFP1 proteins on budding yeast *Saccharomyces cerevisiae* (Sc) and the maize pathogen *Colletotrichum graminicola* (CgM2), which increases virulence in AFP1-silenced plants^8^. We visualized AFP1 fluorescence on the conidial cell poles of CgM2, and the bud sites and the cell division zones of budding yeast, whereas AFP1*-His fluorescence was not detected in these regions (Fig. 6A). While yeast Sc*cda1 and cda2* genes are expressed and required for spore wall assembly^35, 36^, six out of the seven *cda* genes were expressed in the CgM2 conidial cells (Fig. S3B). In the Y2H assay, AFP1 interacted with two of the most highly expressed CgM2 CDAs (GLRG7915 and GLRG386) and with a yeast ScCda2 (Fig. 6B and 6C). The binding of the AFP1 to budding yeast cells resulted in reduced cell viability, as evidenced by a significant decrease in the yeast cell numbers when treated with the AFP1-His proteins. The same reduction in cell viability was not observed in the AFP1*-His-treated yeast cells and mock controls (Fig. 6D). This data is consistent with our previous finding that the AFP1 exhibits antifungal activity against the *U. maydis*^8^. These finding reveals that AFP1 has the ability to inhibit a wide range of fungi by acting on the conserved fungal CDA proteins.

**Fig. 6.**
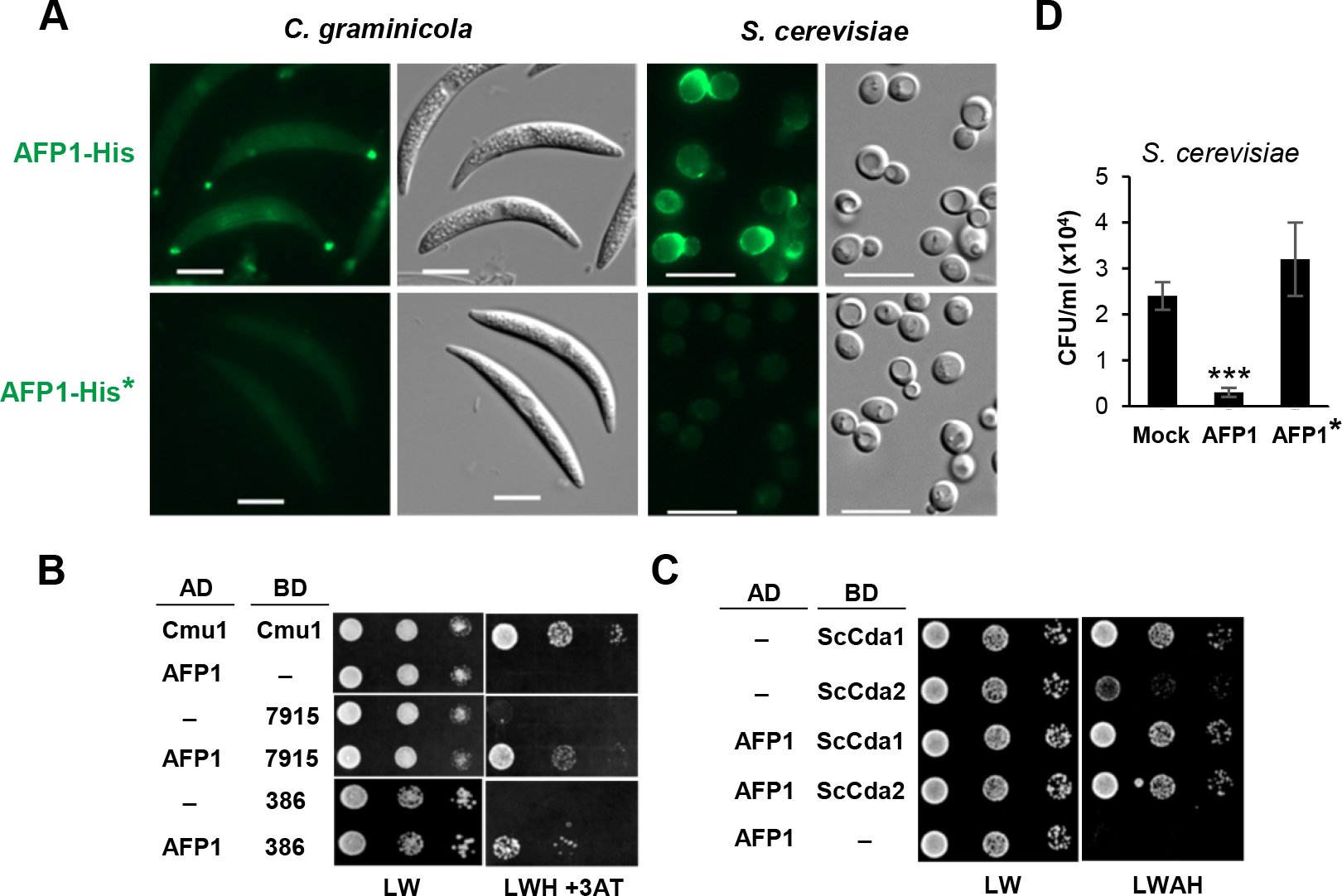
AFP1 acts on CDA of pathogenic and non-pathogenic fungi to affect cell survival. (A) *C. graminicola* and *S. cerevisiae* cells were treated with AFP1-His and AFP1*-His proteins, and followed by immunostaining. Bars:10 μm. (B-C) Yeast two-hybrid assays to detect interaction of AFP1 and CDA proteins of *C. graminicola* (B) or *S. cerevisiae* (C). Yeast transformants containing indicated plasmids were grown on SD/-Leu/-Trp (LW) for growth control and on a SD/-Leu/-Trp/-His/-Ade (LWHA) or LWH plates containing 10 mM of 3AT (3-amino-1,2,4-triazole) to assess protein interaction. Self-interaction of chorismate mutase (Cmu1) was served as positive control, and AD-AFP1/BD and AD/BD-CDA were used as negative controls. Similar results were observed in two independent experiments. Sc: *S. cerevisiae.* (D) *S. cerevisiae* cells incubated with AFP1-His, AFP1*-His and mock (buffer) for 4 hours, serial diluted, and plated on YPAD agars. Colony forming units (CFUs) of yeast cells plating on agar plates were quantified. Values represent mean ± sd of three biological replicates. \*\*\**p* < 0.001 indicates significant differences compared to the CFUs in mock determined by a two-tailed Student’s *t*-test.

## Discussion

In this study, we demonstrate that AFP1 acts on fungal chitin deacetylases (CDAs) to interfere with the conversion of chitin to chitosan. Given the importance and multifaceted roles of CDAs in fungal cell growth across developmental stages and virulence^11, 37–40^, AFP1’s action on the CDAs could be an effective strategy to stop fungal colonization by inhibiting fungal cell growth and spore germination, activating chitin-triggered plant immunity, and ultimately blocking the fungal invasion (Fig. 7). To our knowledge, this is the first report revealing the molecular mechanism of plant CRRSP proteins in plant-pathogen interactions. The maize CRRSP proteins AFP1 displays inhibitory activities against a wide range of fungi, and counteracts the strategy used by *U. maydis* to avoid recognition.

**Fig.7.**
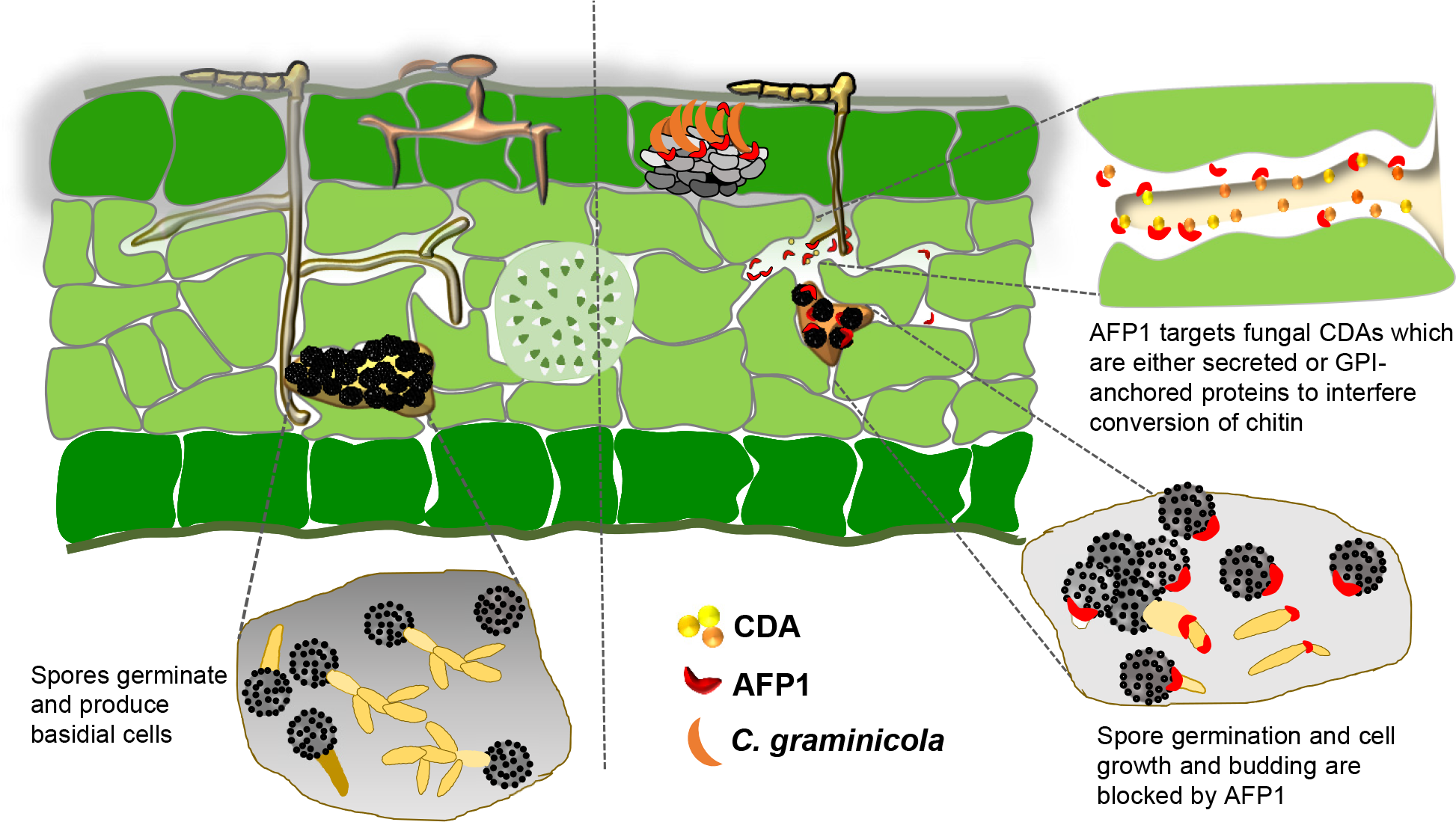
Antifungal action mode of AFP1. Upon pathogen invasion, plants deliver AFP1 proteins to the apoplast to stop the fungal colonization. It interferes with conserved fungal CDA proteins by blocking the conversion of chitin to chitosan, which is necessary for fungal cell development and virulence^11^. By acting on fungal CDAs, AFP1 could prevent spore germination, stop cell budding, and inhibit fungal growth, ultimately leading to fungal cell death and stopping the fungal invasion.

Mannose-binding defective mutant (AFP1*) proteins diminish the binding ability supports that mannose-binding is the prerequisite for the localization of AFP1 on the fungal cell surface. Together with the subsequent finding of a remarkable reduction in AFP1-binding to the bud tips/necks in the Δpmt4 cells where the mannosylated Cda1 and Cda2 localize provide a direct link between mannose-binding property of AFP1 and mannosylated CDA proteins. We envisage that AFP1 could bind to the cell periphery-localized mannosylated CDAs, e.g., Cda4 and Cda2 and GPI-anchored mannosylated CDAs. But, how AFP1 proteins bypass the multilayers of cell walls to reach the GPI-anchored CDAs on the plasma membrane. We anticipated that the CDA-located sites, i.e. cell division, cell poles and hyphal tips are actively growing regions not well covered by cell-wall components, which might expose GPI-anchored CDA proteins and facilitate the access of AFP1 to the targets.

PMT forms a heterodimer or homodimer in *Saccharomyces cerevisiae*^41^. We consider that three PMTs in *Ustilago maydis* likely form a dimer complex to mannosylated their CDA substrates. The smear migration pattern of Cda1/Cda2 in the *pmt4* mutant background could indicate the mannosylation of the proteins by other PMTs (Fig. 4C). PMT1 and PMT2 could also participate in the mannosylation of Cda1, Cda2, and Cda4. At present, we do not know in addition to Cda1 and Cda2 whether other CDAs could also occupy the bud tips/necks and are mannosylated by PMT4. We speculate that Cda7 could be another potential substrate of PMT4 since PMT4 is essential for pathogenesis, and the single deletion of *cda7* reduced fungal virulence ^11^. Although the Cda1/2 localization does not alter in the *pmt4* mutant, we could not conclude that the mannosylation of CDA plays no role in the protein localization. A partial mannosylation in Cda1/2 reduces the interaction with AFP1 but may be sufficient to mediate the protein localization. Due to the essential role of PMT2 in fungal viability^30^, we could not delete all *pmt* genes to study the functionality of mannosylation in CDA localization and understand the PMT-CDA specificity. However, we provide the evidence to support that the mannose-binding of AFP1 and the PMT-dependent mannosylated CDAs are coupled (Fig. 3C and Fig. 4).

In view of the AFP1 interaction with all CDAs in *Ustilago maydis* and other fungal CDAs but not chitinases (CTS) and chitin synthases (CHS) that are putative mannosylated proteins (as predicted by NetOGlyc Server, unpublished analysis), the mannose-binding property of AFP1 appears not to play a role in target selection. The CDA domain of Cda1 is sufficient in mediating the AFP1-Cda1 interaction suggesting that AFP1 likely recognizes the conserved distorted (β/α)8-barrel structure of NodB homologous domain-containing active catalytic sites of CDAs shared by the Carbohydrate Esterase family 4 (CE4) members. According to the phylogenetic analysis of CDAs in the CE4 family, bacterial CDAs and CODs (chitin oligosaccharide deacetylases) are clustered into a different clade and placed at a more related distance to the fungal CDAs^42^. Whether the AFP1 inhibitory could extend to bacterial species will be worth investigating. Based on the observation that AFP1 exhibited the strongest interaction with the Cda1 in the Y2H assay, we reason that AFP1 might have different affinities for each CDA target. The various degrees of mannosylation on CDAs might determine the AFP1-CDA interaction affinity. Alternatively, a specific mannosylation pattern by a different combination of PMT dimers may contribute to the interaction affinity.

Although the interaction of CDA proteins with any fungal membrane-associated proteins has not been reported elsewhere, the localization patterns of CHS^23, 43^ and CDAs are similar in *U. maydis* yeast-like cells. It is tempting to hypothesize that CDAs interact with chitin synthases (CHS) to access de novo nascent chitin substrates and effectively convert them. If this hypothesis holds true, AFP1 targeting on the CDAs might also have an additional negative impact on the interaction between CDAs and CHS, therefore blocking the CDA access to chitin. In the future, it would be interesting to investigate the biological roles of mannosylation on the biophysical properties of CDAs in terms of protein distribution, enzyme activity, and interaction with fungal cell wall proteins.

Our maize AFP1 study provides molecular insights into the understanding of the antifungal mechanisms of plant CRRSP proteins. The finding that AFP1 targets conserved fungal CDAs and displays antagonistic effects against several fungi tested makes its high potential to be developed into an antifungal agent in controlling fungal diseases to reduce threats posed by the fungal kingdom.

## Methods

### Strain construction and growth conditions

The haploid solopathogenic *Ustilago maydis* SG200 strain was used as a reference strain in this study^44, 45^. Strains used and generated in this study are listed in Supplementary Table 1. Plasmid construction using either Gibson Assembly or standard cloning methods as described in Supplementary Table 2. Primers used in each generated plasmid are listed in Supplementary Table 3. A PCR-based approach was used to generate mutants^46^. For gene integration into the *ip* locus or *mig2-6* locus, plasmids containing a carboxin resistant *ip* allele (*ip*R)^47^ or a *mig2-6* allele were linearized with the restriction enzymes SspI or AgeI or EcoN1 and subsequently inserted via homologous recombination. Transformation of *U. maydis* and genomic DNA isolation were performed as described. Positive *U. maydis* transformants were verified by Southern blot analysis.

1. *U. maydis* strains were grown at 28 °C in a liquid YEPSL medium (0.4% yeast extract, 0.4% peptone, and 2% sucrose) and on potato dextrose agar (PDA; 2.4% potato dextrose broth and 2% agar). Yeast cells were grown on YPD agar (BD Difco, USA) at 28 °C and *C. graminicola* CgM2 was grown on oatmeal agar (BD Difco, USA) with continuous exposure to daylight at room temperature.

### Gene expression analysis

SG200 cell pellet was collected after cell grown in liquid YESPL medium reached an OD600 of 0.6. *C. graminicola* (CgM2) conidial cells were harvested from oatmeal agar plates. The cell pellet was washed with sterilized water and frozen in liquid nitrogen. Total RNA extracted using TRIzol (Invitrogen) was subject to DNase-treatment (Promega; Cat#M6101) according to the manufacturer’s recommendation. The cDNA preparation and quantitated RT-PCR analysis were performed as described previously ^8^. The expression of *U. maydis ppi* (peptidyl-prolyl isomerase) was used for normalization. To analyze CgM2 *cda* gene expression using semiquantitative RT-PCR, the *actin* gene expression was used for normalization. PCR amplification was programmed as follows: 95°C for 25 sec, 60°C for 25 sec, and 72°C for 45 sec. After 33 cycles for *cda* amplification and 28 cycles for *actin*, PCR reactions were further incubated at 72°C for 3 min and then chilled at 10°C. Gel images were acquired using the Molecular Imager Gel Doc XR+ system (Bio-RAD) and densitometric analysis was performed using ImageJ^48^.

### Immunolocalization of AFP1 and CDA

The purification of AFP1 proteins from *Nicotiana benthamiana* was performed as described ^8^. To visualize the localization of AFP1-His on yeast-like cells, a final volume of 200 µl of 1x PBS buffer (pH 7.2) containing 4 µg AFP1-His proteins and 1 OD of cells (pre-washed with MES buffer) was incubated for 4 hours at 28 ℃. To visualize AFP1-His localization on germinated spores of FB1xFB2, spores were collected from infected maize leaves and allowed to germinate on PD agar containing 150 μg/ml ampicillin and 35 μg/ml chloramphenicol at 28°C as described^34^. Germinated spores on PD agar were removed, washed, and incubated with 20 µg AFP1-His proteins in a final volume of 200 µl for 4 hours at 28 ℃. The cells/spores were washed with PBS buffer before being subjected to the immunostaining using the primary anti-His antibody and Alexa Fluor 488 (AF488)-conjugated secondary antibody as described^8^.

To visualize the localization of HA-tagged CDA proteins, yeast-like cells were pre-washed before being subjected to immunostaining using primary anti-HA antibody and Alexa Fluor 488 (AF488)-conjugated secondary antibody^8^ in 10mM MES buffer (pH5.5). To visualize the protein localization on filaments of *U. maydis*, cells were induced using hydroxyl fatty acid on parafilm as described^8^ and followed by either an initiated incubation with AFP1-His proteins as described above or directly subjected to immunostaining. AF488 fluorescence was excited at 488 nm and subsequently detected at 500-540 nm using Axio Observer fluorescence microscope equipped with Axiocam 702 Monochrome camera (ZEISS, Germany). Images were processed using ZEN 3.2 imaging software (ZEISS).

### Co-localization and Acceptor photobleaching

The fluorescence-labeled AFP1-His proteins were prepared according to the manufacturer’s instruction of the DyLight 550 Antibody Labeling kit (Thermo Fisher; Cat#84530). One OD600 of cells washed with MES buffer was subjected to immunostaining using the primary anti-HA antibody and AF488-conjugated secondary antibody or wheat germ agglutinin conjugated antibody (WGA-A488; Molecular Probes, Karlsruhe, Germany) to localize CDA proteins or chitin respectively. The immunostained cells or WGA-488 stained cells were washed with MES buffer, followed by adding 4 µg of DyLight 550 labeled AFP1-His proteins (AFP1^550^) to a final volume of 200 µl, and then continued the incubation at room temperature for 4 hours. The AFP1^550^-treated cells were washed and visualized using epi-fluorescence microscopy.

Acceptor photobleaching measurements were performed using a confocal microscope (ZEISS LSM880). The donor (AF488) channel was excited at 488 nm and detected through a 500-540 nm emission filter. The acceptor (DyLight 550) channel was excited at 561 nm and detected through a 580-630 nm emission filter. The images in the donor and acceptor channels were acquired before and following acceptor fluorescent bleaching. Regions of interest (ROI) that overlap with the donor-acceptor fluorescence were selected for photo-destruction by applying 100% laser power at a wavelength of 561 nm for 120 iterations. Evaluation of images was analyzed using the ZEN 3.0 software. The FRET efficiency was calculated according *EFRET = (IDonor(post)-IDonor(pre))/IDonor(post)* where *IDonor(pre)* and *IDonor(post)* are the donor fluorescence intensities prior to and following photo-destruction of the acceptor.

### Live cell time lapse imaging

Spores germinated on PD agar were harvested, incubated with 0.1 µg/µl AFP1-His^550^ proteins in 1x PBS buffer for 4 hours, and mixed with agar to a final concentration of 2% which containing 2% PD medium before being spotted on a chambered coverslip (Ibidi, Catalog #: 80827). Image series were acquired using Olympus DeltaVision Core microscope system with a 63x water immersion objective. Frame time was set at 10 min and a total of 42 frames were acquired. For monitoring AFP1 inhibitory on yeast-like cells, SG200 cells (OD600 = 0.002) were incubated with AFP1-His proteins for 4 hours at room temperature before being spotted on a chambered coverslip to acquire images. Frame time was set at 5 min and a total of 85 frames was acquired in a 7-hour duration.

### CDA activity assay

Cells expressing secreted CDA-Strep proteins were cultured in YEPSL liquid medium until OD600 reached 0.6. A total 75 ml culture supernatant was concentrated and exchanged with buffer W using 10 kD cutoff centrifugal filters (Sartorius, Germany). A 50 µl of Strep-Tactin®XT agarose beads were added to the concentrated sample in a final volume of 1 ml containing 1x protease inhibitor cocktail and incubated overnight at 4 ℃. The beads were washed with 50 mM TEA (Triethanolamine) buffer (pH 8) and aliquoted evenly into three tubes before 10 µg of AFP1-His, AFP1*-His, or TEA buffer was added to a final volume of 350 µl. The incubation was carried out at room temperature for 4 hours and followed by adding GlcNAc5 (A5; Megazyme Cat# O-CHI5) to a final concentration of 0.625 mg/ml and continued the incubation at 37 ℃ for 16 hours. The reaction mixture was centrifuged. The release of acetate in the supernatant was measured by following the manufacturer’s instruction of the Acetic Acid Test Kit (R Biopharm Inc. Cat#10148261035).

### Pull down assay

Cells expressing secreted Strep-tag Cda proteins were grown to OD600 of 0.6 at 28 °C. The 50 ml culture supernatant containing Strep-tagged CDA proteins was concentrated to 1 ml and exchanged to buffer W (100 mM Tris/HCl, 150 mM NaCl, 1 mM EDTA, pH 8) using 10 kD cutoff centrifugal filters (Sartorius, Germany) before being incubated with 25 µl of Strep-Tactin®XT agarose beads (IBA Lifesciences, Germany) overnight at 4 °C in presence of protease inhibitor cocktail (Roche, Switzerland). The beads were washed with buffer W, and incubated with 0.1 µg AFP1-His proteins in a final volume of 500 µl binding buffer (100 mM Tris/HCl, 0.3 M NaCl, 1 mM EDTA, 0.05% NP-40, 1x protease inhibitor cocktail, pH 8) at 4 °C for 2 hours, and then washed with binding buffer. The bound proteins were removed by boiling in sample buffer and subjected to immunoblotting analysis using mouse monoclonal anti-6xHis (Yao-Hong Biotech. Inc., TW) and Strep-tagII (IBA Lifesciences, Germany) primary antibodies and the goat anti-mouse IgG-HRP-conjugated secondary antibody (Thermo Scientific, USA). For the interaction of Cda4-AFP1, 0.2 µg of AFP1-His proteins was incubated with 20 µl Ni-NTA agarose beads (Qiagen, Germany) in buffer (25 mM Tris-Cl, 0.15 M NaCl, 10 mM Imidazole, pH 7.5) for 1 hour at 4 °C. The beads were washed with the same buffer, incubated with the concentrated culture supernatant containing Cda4HA in buffer (25 mM Tris-Cl, 0.3 M NaCl, 0.1% NP-40, 10 mM Imidazole, pH 7.5) at 4 °C for 2 hours. The beads were washed and boiled in a sample buffer to remove the bound proteins for the immunoblotting analysis using mouse monoclonal anti-HA and anti-6xHis antibodies (Yao-Hong Biotech. Inc., TW).

### Yeast two-hybrid assay

Yeast two-hybrid assays were performed as described^49^. Yeast (AH109) transformants containing the desired plasmids were screened on a selective dropout (SD) medium lacking tryptophan (W) and leucine (L) (Clontech). Protein interactions were assessed on SD selection medium lacking LW, histidine (H), and Adenine (A) or lacking LWH and containing 3-amino-1, 2, 4-triazole (3-AT) after three to five days incubation at 28 °C. To detect the expression of proteins in yeast transformants, one OD600 of yeast cells was lysed in a buffer containing 1% β-mercaptoethanol and 0.25 M NaOH. The supernatant fraction was TCA-precipitated and then centrifuged. The protein pellet was dissolved in HU-buffer (100 mM Tris-Cl pH 6.8, 8 M urea, 5% SDS, 0.1 mM EDTA, and 0.1% bromophenol blue) and analyzed by immunoblotting.

### Yeast growth inhibition assay

For the yeast growth inhibition assay, a 100 µl final volume of a reaction containing yeast cells BY4742 (OD600 =0.001) and 20 µg AFP1 proteins were incubated at 28°C for 4 hours in 10mM MES buffer (pH5.5). The cells were serial-diluted, spread on YPD plates (3 plates per each dilution), and incubated at 28°C for two days until colonies appeared. Colony numbers were then manually counted.

### Author contributions

LSM conceived and designed the study and wrote the manuscript. WLT performed immunostaining and FRET analysis. WLT and RMK did live-cell imaging and time-lapse microscopy. RMK performed the initial AFP1 binding analyses. MYX did protein purification. MYX, WLT, and YHL generated constructs and CDA overexpression strains. MYX, WLT, YHL, FAD, and HCL performed Y2H assays. YHL analyzed *cda* gene expression. WLT and LSM did pull-down assays. WLT and FAD analyzed CDA activity inhibition assay and yeast growth-inhibition assay. All authors discussed the results and commented on the manuscript.

## Acknowledgements

We are indebted to Regine Kahmann for supply of the *cda* deletion mutants and other strains (Import permit# 109-F-500 and 107-F-002) and critically reading the manuscript. We are grateful to Erh-Min Lai, Chih-Hang Wu, and Ooi-Kock Teh for suggestions and comments on the manuscript. We thank Prof. Dr. Bruno Moerschbacher and Mounashree J. Urs for providing technical advice on CDA activity analysis. We acknowledge Ji-Ying Huang in the Plant Cell Biology Core Laboratory for assisting in live-cell imaging and FRET microscopy and the staff in the DNA Sequencing Core Laboratory at the Institute of Plant and Microbial Biology for technical assistance. We thank Chibbhi K. Bhaskar and Cuong V. Hoang for their comments on an early version of the manuscript. Our work is supported by funds from the Ministry of Science and Technology, Taiwan (MOST# 107-2311-B-001 -044 -MY3).

## Supplementary Information

**Fig. S1.**
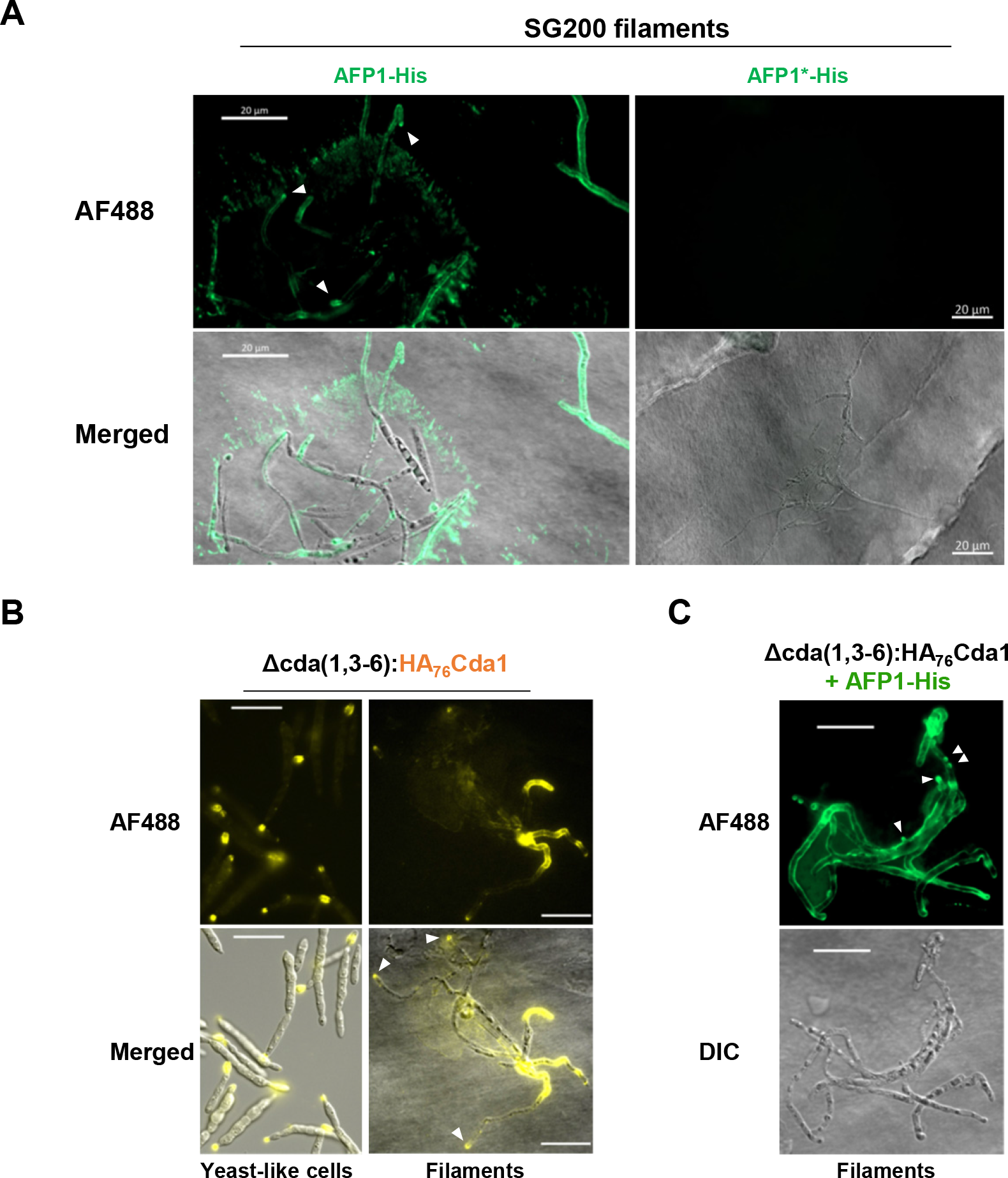
Immunostaining of AFP1 and Cda1 in *U. maydis* cells. (A) Filaments of SG200 induced by hydroxyl fatty acids on hydrophobic surface were treated with AFP1-His and AFP1*-His proteins, and followed by immunostaining with an anti-His antibody and an AF488-conjugated secondary antibody. Bars, 20 μm. (B) Yeast-like cells and filaments of Δcda(1,3-6):HA_76_Cda1 strain which constitutively expressed HA_76_Cda1 proteins under promoter *otef* in Δcda(1,3-6) deletion mutant were subject to immunostaining using an anti-HA antibody and an AF488-conjugated secondary antibody. (C) Filaments of Δcda(1,3-6):HA_76_Cda1 strain treated with AFP1-His proteins before subjected to immunostaining using an anti-His antibody and an AF488-conjugated secondary antibody. Bars, 20 μm. Arrow heads indicate AFP1-His or HA_76_Cda1 fluorescence on hyphal tips.

**Fig. S2.**
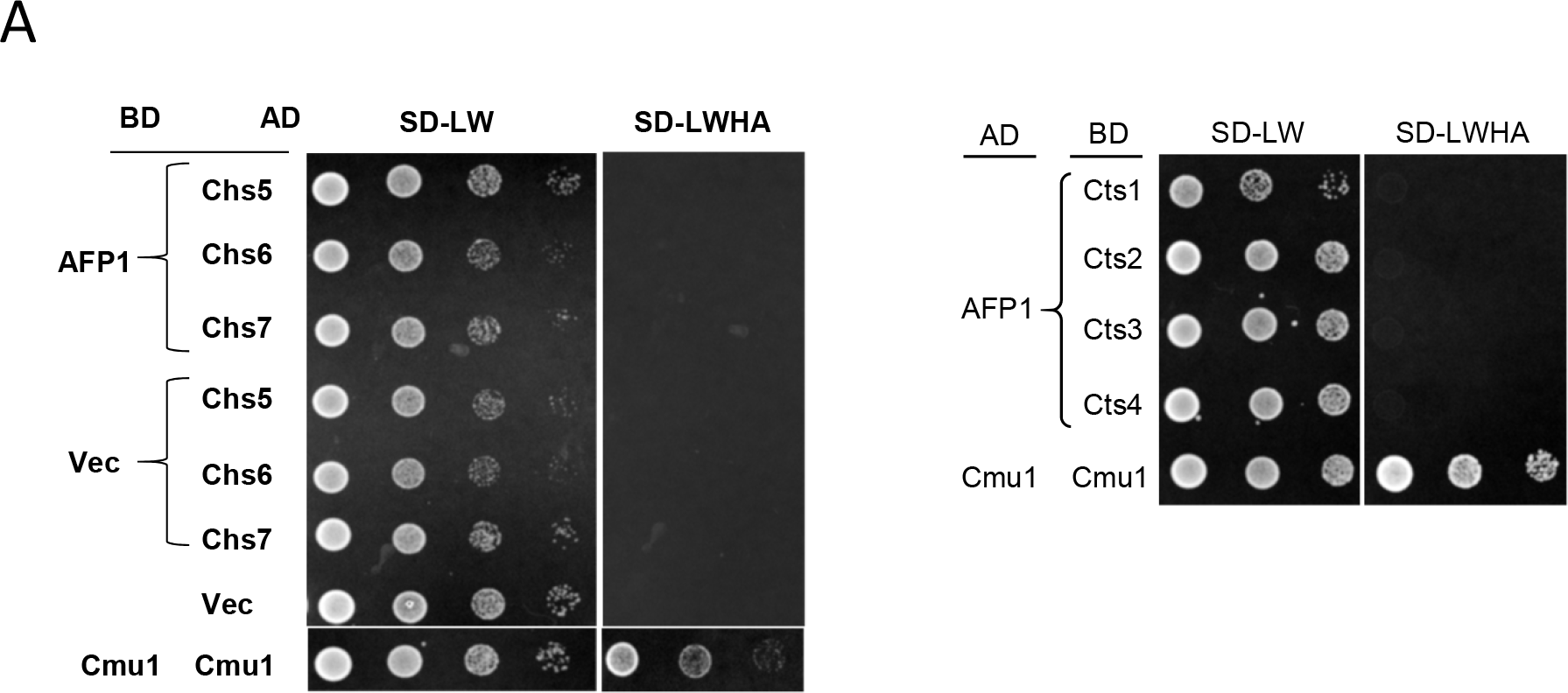
Yeast-two hybrid analysis of AFP1-CHS (chitin synthases) and AFP1-CTS (chitinase) interactions. Yeast two-hybrid assay to detect interaction of AFP1 and CHS/CTS proteins. Yeast cells containing the indicated plasmids were grown on SD/-Leu/-Trp (LW) and SD/-Leu/-Trp/-His/-Ade (LWHA) plates for 2-3 days. Self-interaction of chorismate mutase (Cmu1) was served as the positive control and AD-AFP1/BD and BD-AFP1/AD and BD/CHS as negative controls. Similar results were observed in at least two independent experiments.

**Fig. S3.**
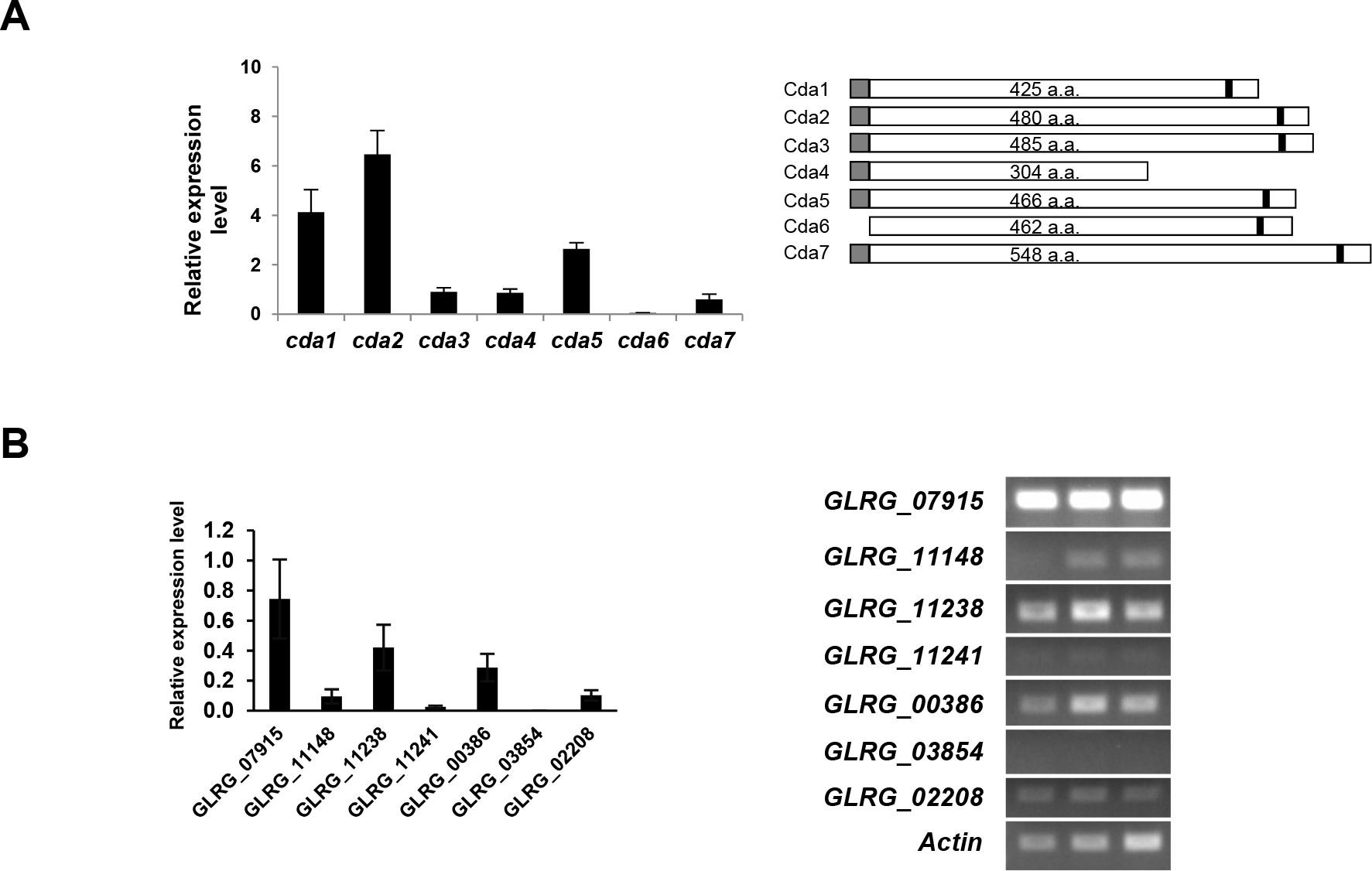
*cda* gene expression of *U. maydis and C. graminicola*. (A) Total RNA was extracted rom SG200 cells grown in YEPSL liquid medium and subjected to quantitative RT-PCR. xpression levels of *U. maydis cda* gene were normalized relative to the constitutively xpressed peptidyl-prolyl isomerase (ppi). Three biological replicates were analyzed. Values epresent mean ± sd. Right panel showed the schematic drawings of CDA proteins with ndicated signal peptides (grey boxes) and predicted GPI anchors (black boxes). a. a., amino cids. (B) Total RNA was extracted from *C. graminicola* conidial cells grown on agar plates and ubjected to semi-quantitative RT-PCR. Values were obtained by quantified intensities of PCR ands in the right panel using ImageJ software, and were normalized relative to the onstitutively expressed *actin* gene. Data represent mean ± sd of the three biological replicates hown in right panel.

**Fig. S4.**
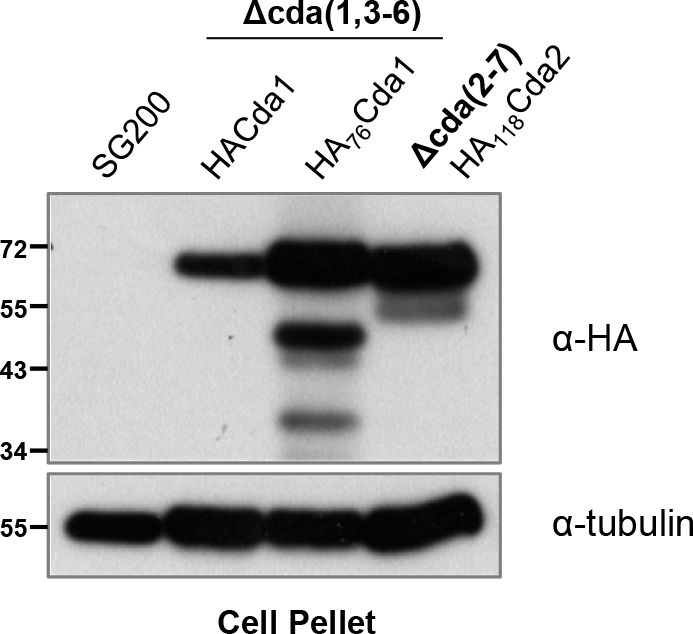
Immunoblot analysis of HA-tagged CDA protein expression. HA-tagged CDA proteins ere expressed under constitutive promoter *otef* in the indicated deletion mutants. The ndicated strains grown in YEPSL liquid medium to OD_600_ of 0.6-0.7 were harvested and djusted to OD_600_ of 20. Proteins from cell pellets were prepared and subjected to immunoblot nalysis using anti-HA or anti tubulin antibodies as indicated.

**Fig. S5.**
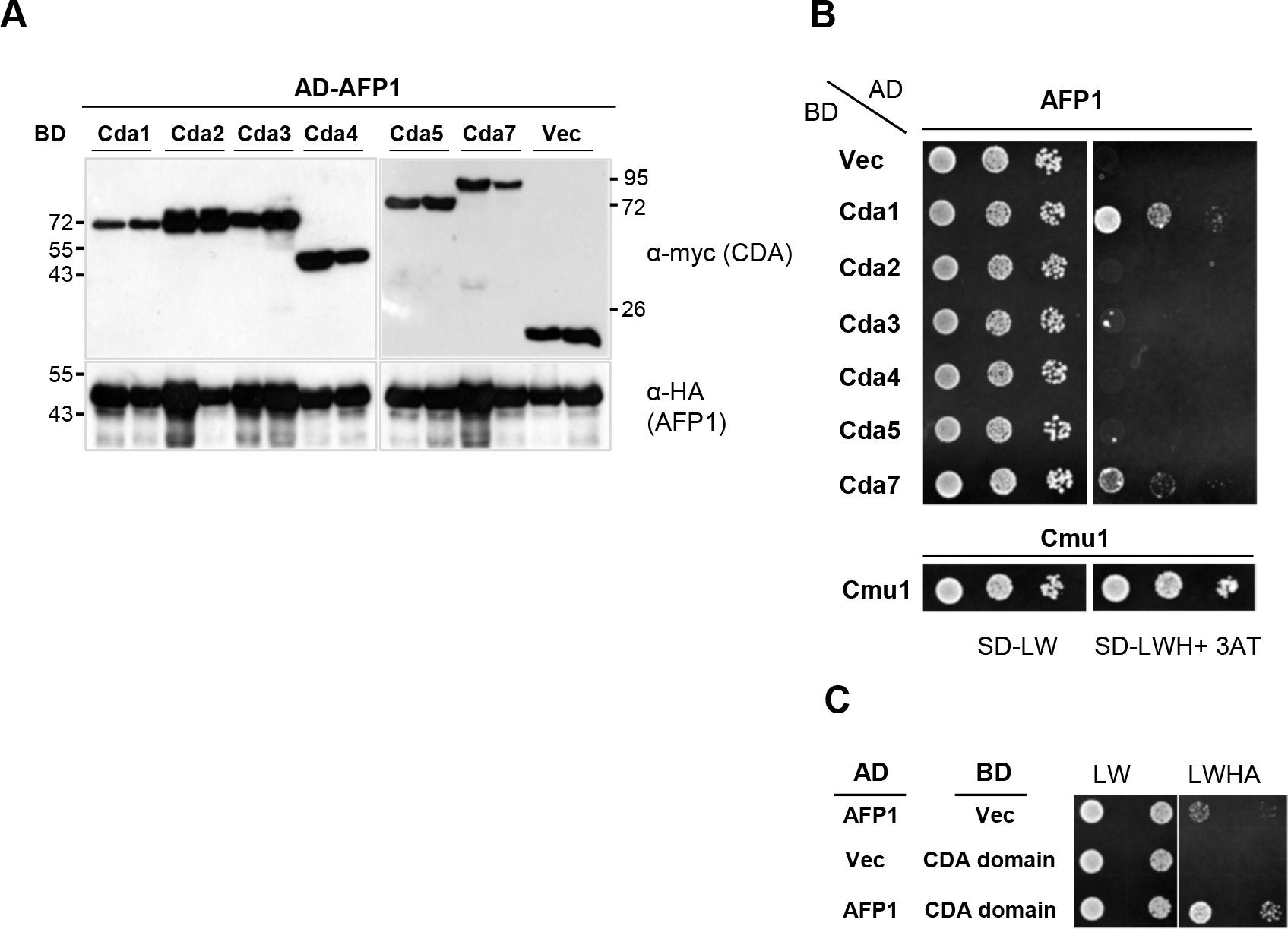
Yeast-two hybrid analysis of AFP1-CDA interaction. (A) Immunoblot analysis of CDA and AFP1 protein expression in yeast transformants. Yeast cells containing indicated plasmids were grown in YPD liquid medium until OD_600_ of 0.7-0.8. One OD of cell pellet was collected, lysed, and TCA precipitated and analyzed by immunoblots using indicated antibodies. Two independent clones were selected for analysis. (B, C) Y2H assay to detect interaction of AFP1 with (B) CDA proteins and (C) CDA domain of Cda1 protein. Yeast cells with indicated plasmids were grown on SD-LW and SD-LWHA or SD-LWH containing 1mM of 3AT plates for 3 days. Self-interaction of chorismate mutase (Cmu1) was served as the positive control. Similar results were observed in at least two independent experiments.

**Fig. S6.**
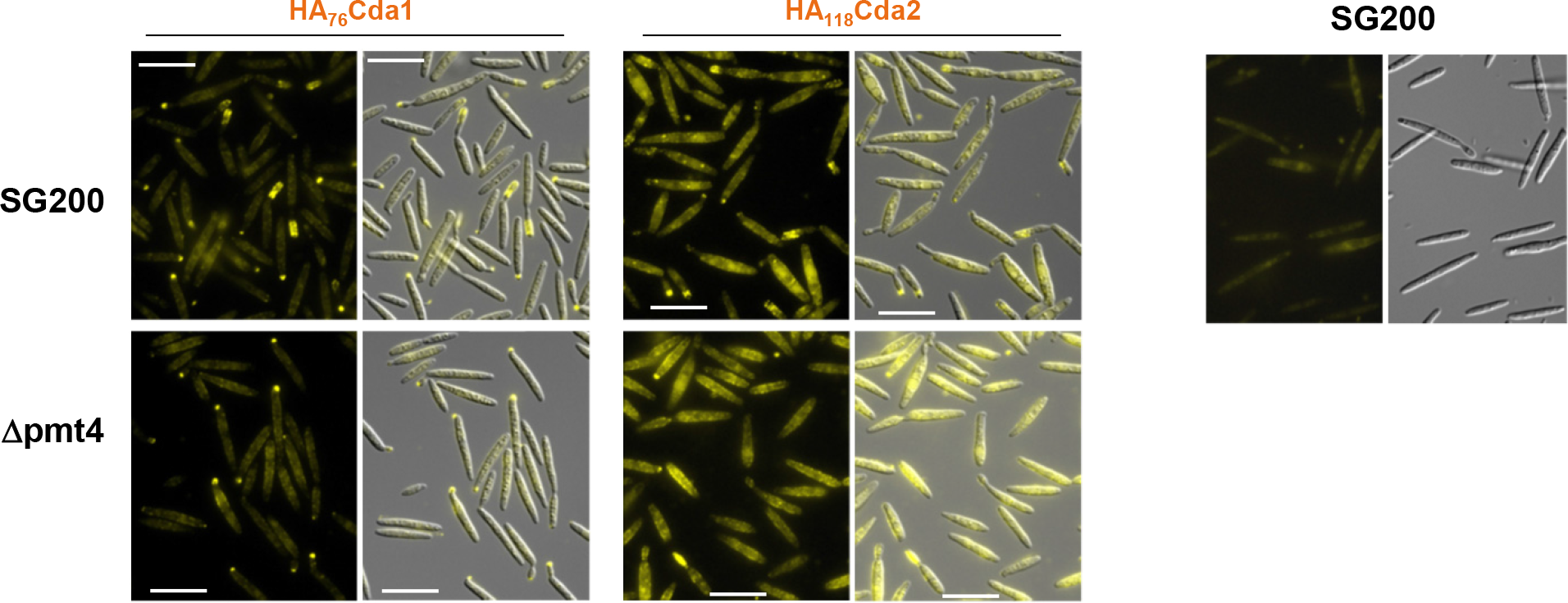
Immunolocalization of HA_76_Cda and HA_118_Cda2. Cells expressing HA-tagged CDA proteins under constitutive promoter *otef* in indicated strains were subjected to immunostaining using anti-HA antibody and an AF488-conjugated secondary antibody to localize CDA proteins. Bars, 20 μm. (Right) Wild-type SG200 cells subjected to immunostaining using anti-HA antibody and AF488-conjugated secondary antibody was served as negative control.

**Fig. S7.**
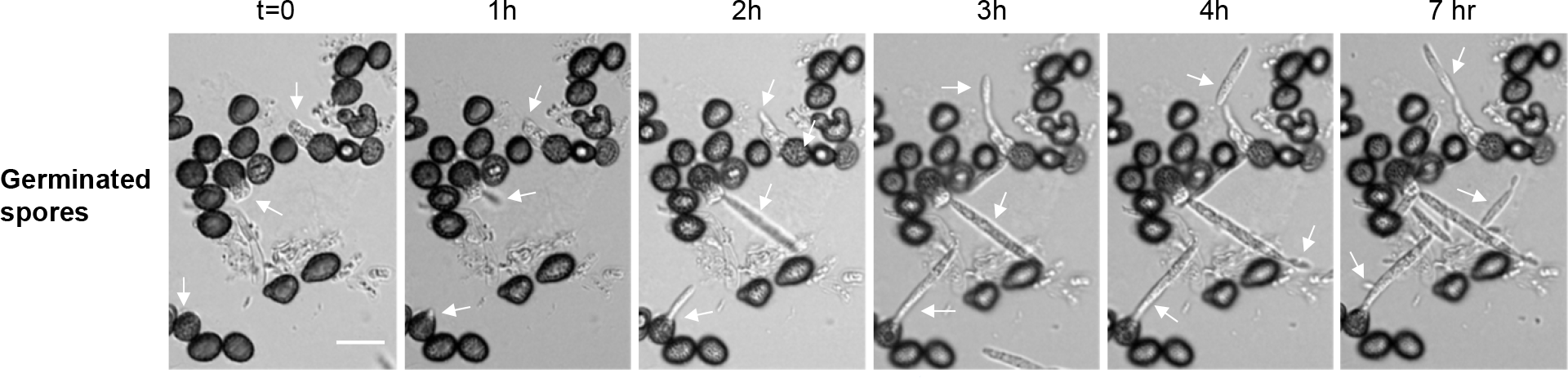
Spore germination in absence of AFP1 proteins. *U. maydis* FB1x FB2 germinated spores in agar containing 2% PD medium. Bars,10 μm. Images of germinated spores without AFP1 proteins added were acquired at indicated time points. White arrows track emergence of promycelium over incubation time.

**Supplementary Table 1:**
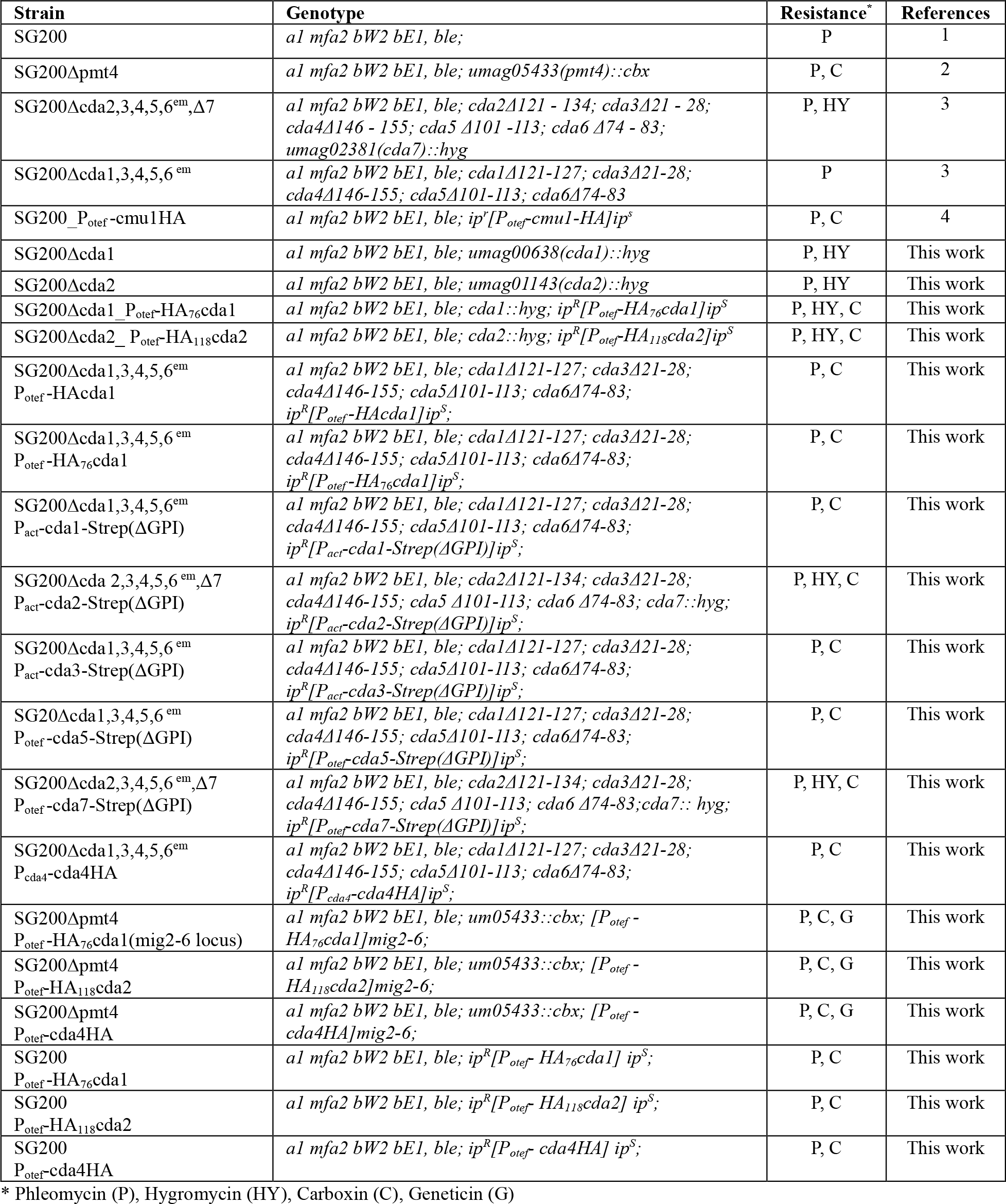
Strains used in this study

**Supplementary Table 2:**
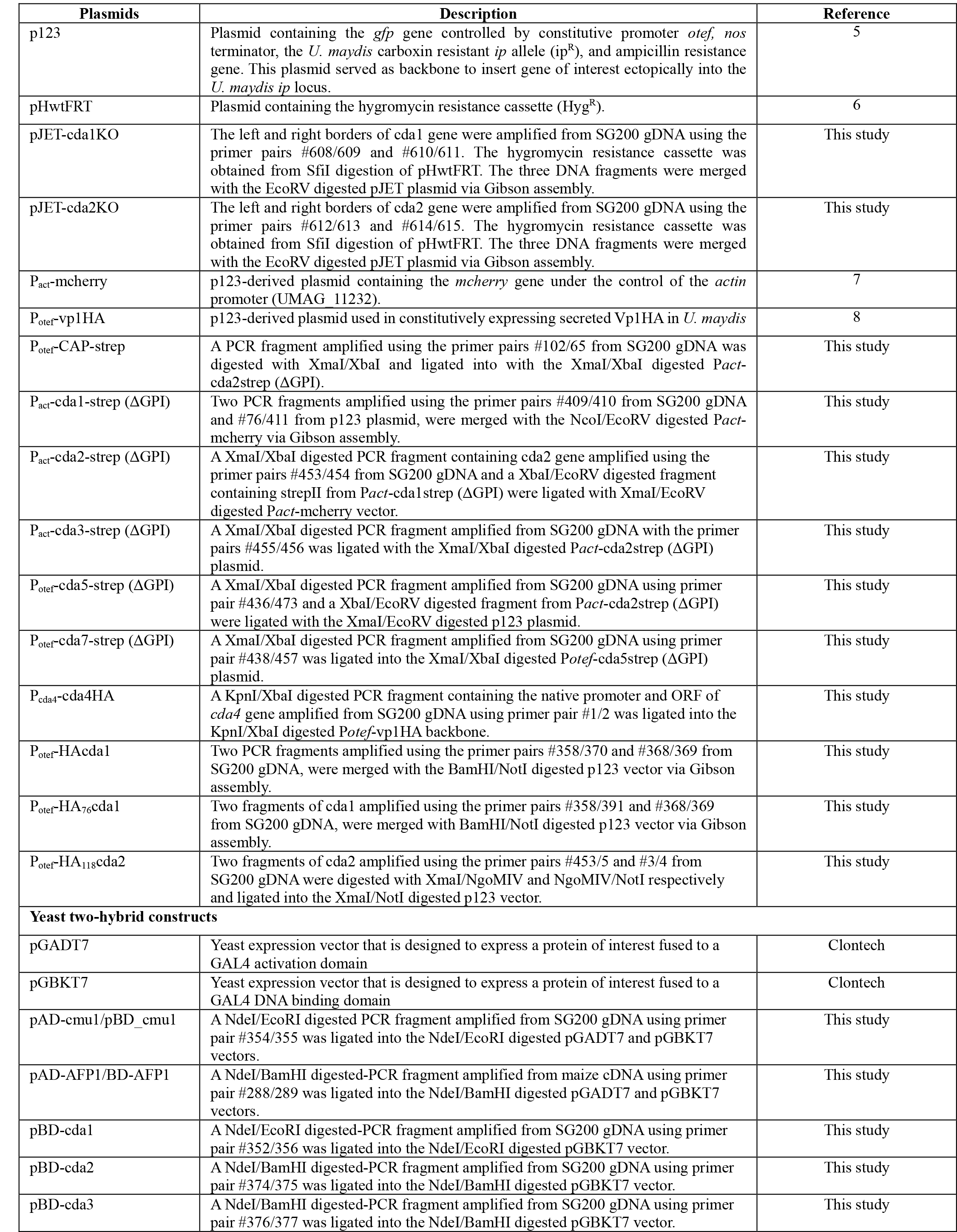
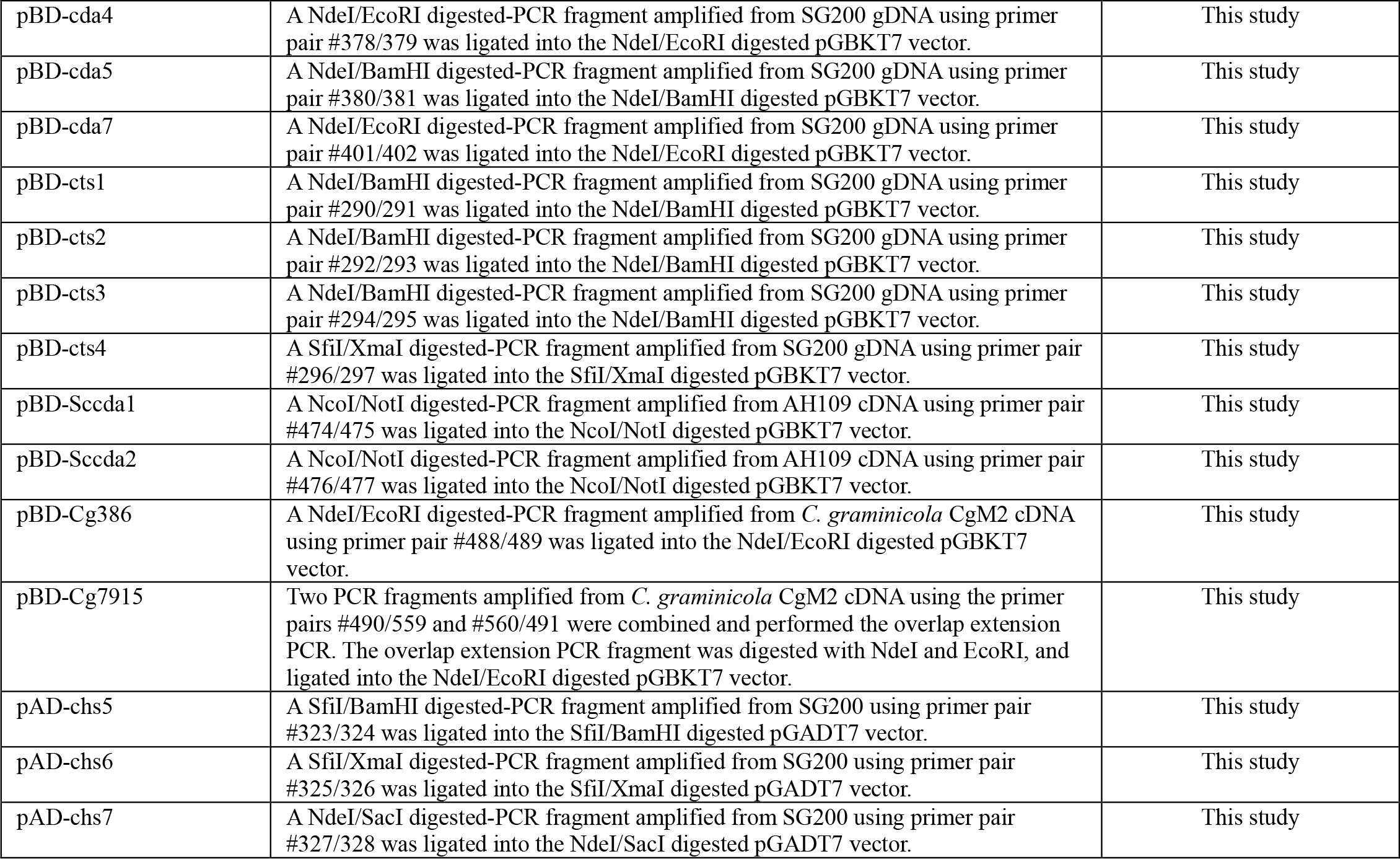
**Plasmids used in this study**

**Supplementary Table 3:**
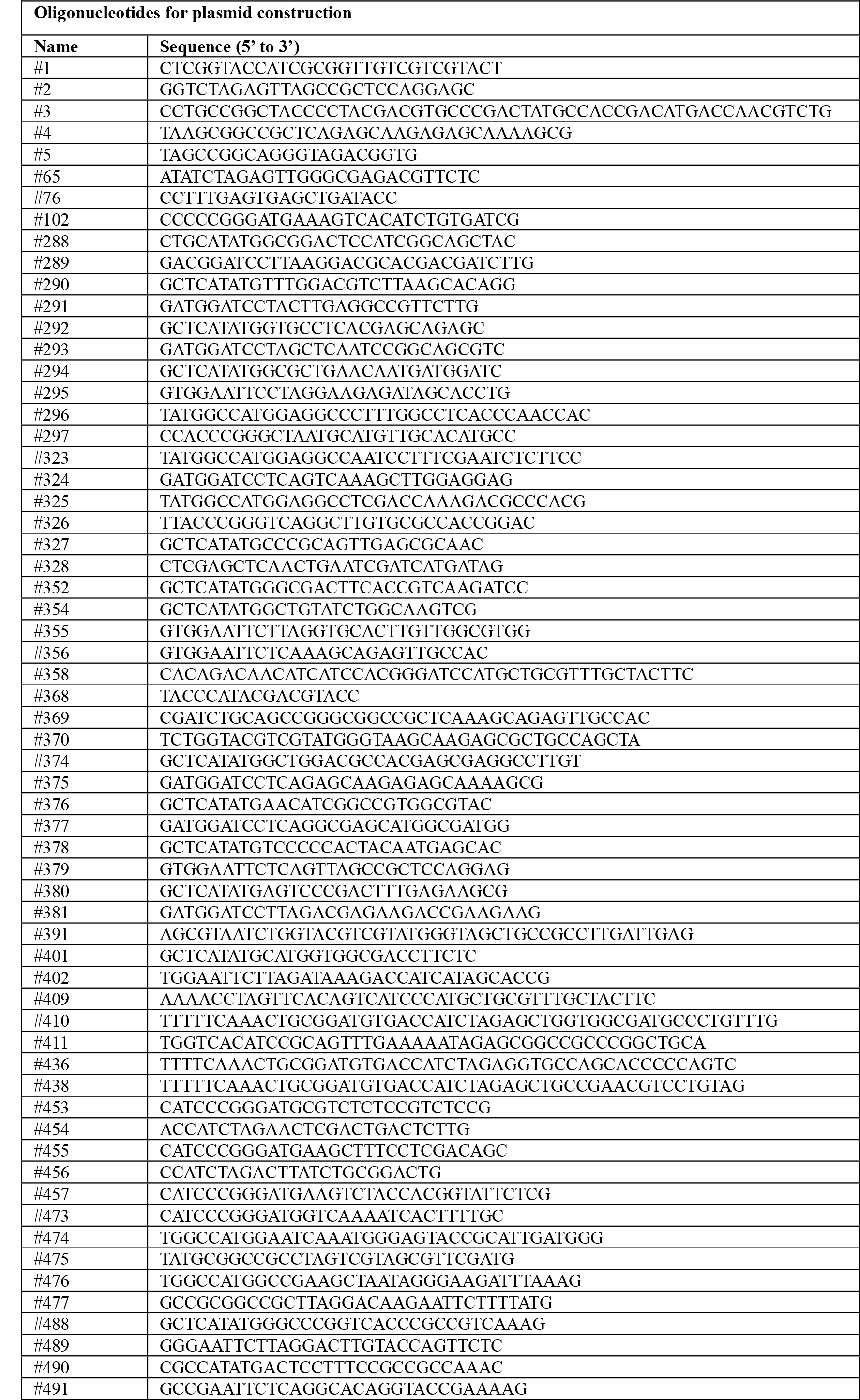
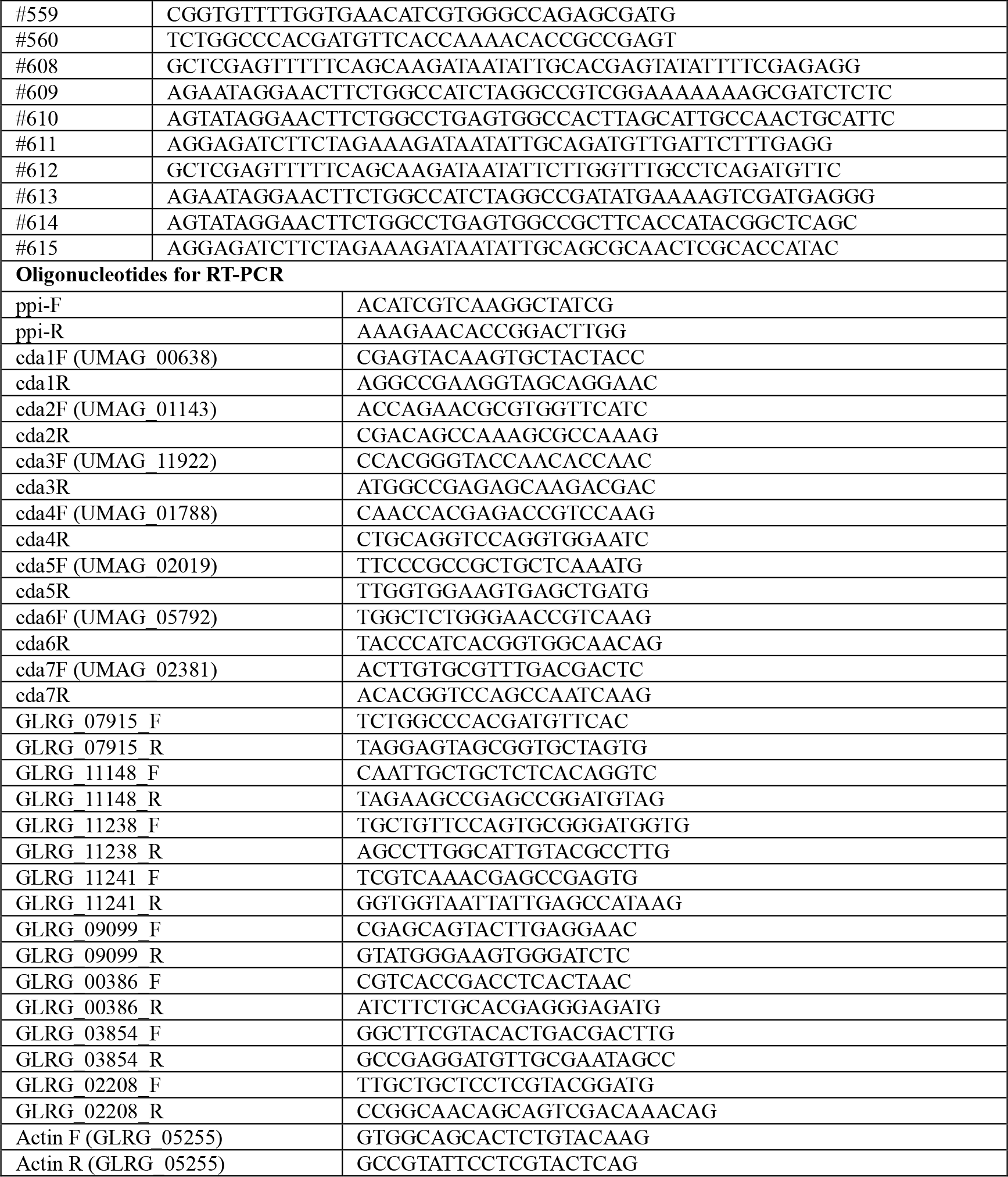
**Oligonucleotides used in this study**

## References

1. Jones, J. D. & Dangl, J. L. The plant immune system. Nature 444, 323–329, doi:10.1038/nature05286 (2006).

2. Monaghan, J. & Zipfel, C. Plant pattern recognition receptor complexes at the plasma membrane. Curr Opin Plant Biol 15, 349–357, doi:10.1016/j.pbi.2012.05.006 (2012).

3. Doehlemann, G. & Hemetsberger, C. Apoplastic immunity and its suppression by filamentous plant pathogens. New Phytol 198, 1001–1016, doi:10.1111/nph.12277 (2013).

4. Buscaill, P. & van der Hoorn, R. A. L. Defeated by the nines: nine extracellular strategies to avoid MAMP recognition in plants. Plant Cell, doi:10.1093/plcell/koab109 (2021).

5. Lanver, D. et al. Ustilago maydis effectors and their impact on virulence. Nat Rev Microbiol 15, 409–421, doi:10.1038/nrmicro.2017.33 (2017).

6. Lanver, D. et al. The Biotrophic Development of Ustilago maydis Studied by RNA-Seq Analysis. Plant Cell 30, 300–323, doi:10.1105/tpc.17.00764 (2018).

7. Okmen, B. et al. Dual function of a secreted fungalysin metalloprotease in Ustilago maydis. New Phytol 220, 249–261, doi:10.1111/nph.15265 (2018).

8. Ma, L. S. et al. The Ustilago maydis repetitive effector Rsp3 blocks the antifungal activity of mannose-binding maize proteins. Nat Commun 9, 1711, doi:10.1038/s41467-018-04149-0 (2018).

9. Mueller, A. N., Ziemann, S., Treitschke, S., Assmann, D. & Doehlemann, G. Compatibility in the Ustilago maydis-maize interaction requires inhibition of host cysteine proteases by the fungal effector Pit2. PLoS Pathog 9, e1003177, doi:10.1371/journal.ppat.1003177 (2013).

10. Hemetsberger, C., Herrberger, C., Zechmann, B., Hillmer, M. & Doehlemann, G. The Ustilago maydis effector Pep1 suppresses plant immunity by inhibition of host peroxidase activity. PLoS Pathog 8, e1002684, doi:10.1371/journal.ppat.1002684 (2012).

11. Rizzi, Y. S. et al. Chitosan and Chitin Deacetylase Activity Are Necessary for Development and Virulence of Ustilago maydis. mBio 12, doi:10.1128/mBio.03419-20 (2021).

12. Chen, Z. A superfamily of proteins with novel cysteine-rich repeats. Plant Physiol 126, 473–476 (2001).

13. Vaattovaara, A. et al. Mechanistic insights into the evolution of DUF26-containing proteins in land plants. Commun Biol 2, 56, doi:10.1038/s42003-019-0306-9 (2019).

14. Czernic, P. et al. Characterization of an Arabidopsis thaliana receptor-like protein kinase gene activated by oxidative stress and pathogen attack. Plant J 18, 321–327 (1999).

15. Yadeta, K. A. et al. A Cysteine-Rich Protein Kinase Associates with a Membrane Immune Complex and the Cysteine Residues Are Required for Cell Death. Plant Physiol 173, 771–787, doi:10.1104/pp.16.01404 (2017).

16. Ohtake, Y., Takahashi, T. & Komeda, Y. Salicylic acid induces the expression of a number of receptor-like kinase genes in Arabidopsis thaliana. Plant Cell Physiol 41, 1038–1044 (2000).

17. Hunter, K. et al. CRK2 Enhances Salt Tolerance by Regulating Callose Deposition in Connection with PLDalpha1. Plant Physiol 180, 2004–2021, doi:10.1104/pp.19.00560 (2019).

18. Lee, J. Y. et al. A plasmodesmata-localized protein mediates crosstalk between cell-to-cell communication and innate immunity in Arabidopsis. Plant Cell 23, 3353–3373, doi:10.1105/tpc.111.087742 (2011).

19. Kim, S. G. et al. In-depth insight into in vivo apoplastic secretome of rice-Magnaporthe oryzae interaction. J Proteomics 78, 58–71, doi:10.1016/j.jprot.2012.10.029 (2013).

20. Miyakawa, T. et al. A secreted protein with plant-specific cysteine-rich motif functions as a mannose-binding lectin that exhibits antifungal activity. Plant Physiol 166, 766–778, doi:10.1104/pp.114.242636 (2014).

21. Han, L. B. et al. The Cotton Apoplastic Protein CRR1 Stabilizes Chitinase 28 to Facilitate Defense against the Fungal Pathogen Verticillium dahliae. Plant Cell 31, 520–536, doi:10.1105/tpc.18.00390 (2019).

22. Treitschke, S., Doehlemann, G., Schuster, M. & Steinberg, G. The myosin motor domain of fungal chitin synthase V is dispensable for vesicle motility but required for virulence of the maize pathogen Ustilago maydis. Plant Cell 22, 2476–2494, doi:10.1105/tpc.110.075028 (2010).

23. Weber, I., Assmann, D., Thines, E. & Steinberg, G. Polar localizing class V myosin chitin synthases are essential during early plant infection in the plant pathogenic fungus Ustilago maydis. Plant Cell 18, 225–242, doi:10.1105/tpc.105.037341 (2006).

24. Langner, T. et al. Chitinases Are Essential for Cell Separation in Ustilago maydis. Eukaryot Cell 14, 846–857, doi:10.1128/EC.00022-15 (2015).

25. Kenworthy, A. K. Imaging protein-protein interactions using fluorescence resonance energy transfer microscopy Methods 24, 289–296, doi:10.1006/meth.2001.1189 (2001).

26. Kenworthy, A. K., Petranova, N. & Edidin, M. High-resolution FRET microscopy of cholera toxin B-subunit and GPI-anchored proteins in cell plasma membranes. Mol Biol Cell 11, 1645–1655, doi:10.1091/mbc.11.5.1645 (2000).

27. Karpova, T. S. et al. Fluorescence resonance energy transfer from cyan to yellow fluorescent protein detected by acceptor photobleaching using confocal microscopy and a single laser. J Microsc 209, 56–70, doi:10.1046/j.1365-2818.2003.01100.x (2003).

28. Djamei, A. et al. Metabolic priming by a secreted fungal effector. Nature 478, 395–398, doi:10.1038/nature10454 (2011).

29. Zhao, Y., Park, R. D. & Muzzarelli, R. A. Chitin deacetylases: properties and applications. Mar Drugs 8, 24–46, doi:10.3390/md8010024 (2010).

30. Fernandez-Alvarez, A., Elias-Villalobos, A. & Ibeas, J. I. The O-mannosyltransferase PMT4 is essential for normal appressorium formation and penetration in Ustilag maydis. Plant Cell 21, 3397–3412, doi:10.1105/tpc.109.065839 (2009).

31. Banuett, F. & Herskowitz, I. Discrete developmental stages during teliospore formation in the corn smut fungus, Ustilago maydis. Development 122, 2965–2976 (1996).

32. Sacadura, N. T. & Saville, B. J. Gene expression and EST analyses of Ustilago maydis germinating teliospores. Fungal Genet Biol 40, 47–64, doi:10.1016/s1087-1845(03)00078-1 (2003).

33. Ostrowski, L. A., Seto, A. M. & Saville, B. Investigating Teliospore Germination Using Microrespiration Analysis and Microdissection. J Vis Exp, doi:10.3791/57628 (2018).

34. Fukada, F., Rossel, N., Munch, K., Glatter, T. & Kahmann, R. A small Ustilago maydis effector acts as a novel adhesin for hyphal aggregation in plant tumors. New Phytol 231, 416–431, doi:10.1111/nph.17389 (2021).

35. Christodoulidou, A., Briza, P., Ellinger, A. & Bouriotis, V. Yeast ascospore wall assembly requires two chitin deacetylase isozymes. FEBS Lett 460, 275–279, doi:10.1016/s0014-5793(99)01334-4 (1999).

36. Christodoulidou, A., Bouriotis, V. & Thireos, G. Two sporulation-specific chitin deacetylase-encoding genes are required for the ascospore wall rigidity of Saccharomyces cerevisiae. J Biol Chem 271, 31420–31425, doi:10.1074/jbc.271.49.31420 (1996).

37. Upadhya, R. et al. Cryptococcus neoformans Cda1 and Its Chitin Deacetylase Activity Are Required for Fungal Pathogenesis. mBio 9, doi:10.1128/mBio.02087-18 (2018).

38. Gao, F. et al. Deacetylation of chitin oligomers increases virulence in soil-borne fungal pathogens. Nat Plants 5, 1167–1176, doi:10.1038/s41477-019-0527-4 (2019).

39. Geoghegan, I. A. & Gurr, S. J. Chitosan Mediates Germling Adhesion in Magnaporthe oryzae and Is Required for Surface Sensing and Germling Morphogenesis. PLoS Pathog 12, e1005703, doi:10.1371/journal.ppat.1005703 (2016).

40. Baker, L. G., Specht, C. A., Donlin, M. J. & Lodge, J. K. Chitosan, the deacetylated form of chitin, is necessary for cell wall integrity in Cryptococcus neoformans. Eukaryot Cell 6, 855–867, doi:10.1128/EC.00399-06 (2007).

41. Girrbach, V. & Strahl, S. Members of the evolutionarily conserved PMT family of protein O-mannosyltransferases form distinct protein complexes among themselves. J Biol Chem 278, 12554–12562, doi:10.1074/jbc.M212582200 (2003).

42. Grifoll-Romero, L., Pascual, S., Aragunde, H., Biarnes, X. & Planas, A. Chitin Deacetylases: Structures, Specificities, and Biotech Applications. Polymers (Basel) 10, doi:10.3390/polym10040352 (2018).

43. Weber, I., Gruber, C. & Steinberg, G. A class-V myosin required for mating, hyphal growth, and pathogenicity in the dimorphic plant pathogen Ustilago maydis. Plant Cell 15, 2826–2842, doi:10.1105/tpc.016246 (2003).

44. Muller, P., Aichinger, C., Feldbrugge, M. & Kahmann, R. The MAP kinase kpp2 regulates mating and pathogenic development in Ustilago maydis. Mol Microbiol 34, 1007–1017, doi:10.1046/j.1365-2958.1999.01661.x (1999).

45. Kamper, J. et al. Insights from the genome of the biotrophic fungal plant pathogen Ustilago maydis. Nature 444, 97–101, doi:10.1038/nature05248 (2006).

46. Kamper, J. A PCR-based system for highly efficient generation of gene replacement mutants in Ustilago maydis. Mol Genet Genomics 271, 103–110, doi:10.1007/s00438-003-0962-8 (2004).

47. Broomfield, P. L. & Hargreaves, J. A. A single amino-acid change in the iron-sulphur protein subunit of succinate dehydrogenase confers resistance to carboxin in Ustilago maydis. Curr Genet 22, 117–121, doi:10.1007/BF00351470 (1992).

48. Schneider, C. A., Rasband, W. S. & Eliceiri, K. W. NIH Image to ImageJ: 25 years of image analysis. Nat Methods 9, 671–675, doi:10.1038/nmeth.2089 (2012).

49. Tanaka, S. et al. A secreted Ustilago maydis effector promotes virulence by targeting anthocyanin biosynthesis in maize. Elife 3, e01355, doi:10.7554/eLife.01355 (2014).

## References

1. Kamper, J. et al. Insights from the genome of the biotrophic fungal plant pathogen Ustilago maydis. Nature 444, 97-101, doi:10.1038/nature05248 (2006).

2. Fernandez-Alvarez, A., Elias-Villalobos, A. & Ibeas, J. I. The O-mannosyltransferase PMT4 is essential for normal appressorium formation and penetration in Ustilago maydis. Plant Cell 21, 3397–3412, doi:10.1105/tpc.109.065839 (2009).

3. Rizzi, Y. S. et al. Chitosan and Chitin Deacetylase Activity Are Necessary for Development and Virulence of Ustilago maydis. mBio 12, doi:10.1128/mBio.03419-20 (2021).

4. Djamei, A. et al. Metabolic priming by a secreted fungal effector. Nature 478, 395–398, doi:10.1038/nature10454 (2011).

5. Aichinger, C. et al. Identification of plant-regulated genes in Ustilago maydis by enhancer-trapping mutagenesis. Mol Genet Genomics 270, 303–314, doi:10.1007/s00438-003-0926-z (2003).

6. Khrunyk, Y., Munch, K., Schipper, K., Lupas, A. N. & Kahmann, R. The use of FLP-mediated recombination for the functional analysis of an effector gene family in the biotrophic smut fungus Ustilago maydis. New Phytol 187, 957–968, doi:10.1111/j.1469-8137.2010.03413.x (2010).

7. Lanver, D. et al. The Biotrophic Development of Ustilago maydis Studied by RNA-Seq Analysis. Plant Cell 30, 300–323, doi:10.1105/tpc.17.00764 (2018).

8. Cuong V. Hoang, C. K. B., Lay-Sun Ma. A Novel Core Effector Vp1 Promotes Fungal Colonization and Virulence of *Ustilago maydis*. J. of Fungi 7, doi:10.3390/jof7080589 (2021).

